# Inverse stable isotope labeling (InverSIL) links predicted catecholate siderophore gene clusters to their products in diverse bacteria

**DOI:** 10.1101/2025.11.05.686764

**Authors:** Jose Miguel D. Robes, Tashi C. E. Liebergesell, Victoria P. Medvedeva, Aaron W. Puri

## Abstract

Bacteria produce high-affinity iron-chelating secondary metabolites called siderophores to access insoluble Fe(III) in their environments. Genome mining has revealed many predicted siderophore biosynthetic gene clusters (BGCs) in bacterial genomes, however the structures of their siderophore products remain mostly undetermined. This limits our molecular-level understanding of how bacteria acquire iron, as well as how they interact with other taxa that may use the same siderophores within bacterial communities. Here, we apply inverse stable isotope labeling (InverSIL) to rapidly connect predicted siderophore BGCs to their products. With InverSIL, bacteria are grown on ^13^C-substituted carbon sources and then fed predicted biosynthetic precursors at their natural isotopic abundance to identify BGC products by mass spectrometry, which removes issues with the availability of isotopically substituted precursors. We use InverSIL to determine the structures of the siderophore products of predicted BGCs from the methylotrophic genera *Methylophilus* and *Methylobacterium*, and the siderophores produced by the opportunistic pathogen *Chromobacterium violaceum*, which were previously shown to be essential for virulence yet remained structurally uncharacterized. We next use this approach to reveal the unexpected production of enterobactin by the genera *Kushneria* and *Paracoccus*, which was difficult to predict from genome sequences due to the distributed nature of the biosynthetic genes within the genomes. Finally, we use InverSIL to discover new siderophores, cellulochelin A and B, from the cellulose-degrading plant symbiont *Cellulomonas* sp. strain Leaf334. These findings demonstrate the utility of InverSIL for functional BGC characterization and expand our molecular understanding of bacterial iron acquisition strategies.

**IMPORTANCE:** Iron acquisition is essential for microbial survival, and bacteria produce secondary metabolites called siderophores to scavenge iron from the environment. While bacterial genome sequences show many predicted genes for making siderophores, most remain unlinked to their metabolic products. Understanding which siderophores bacteria produce is critical for elucidating microbial iron acquisition strategies, ecological interactions, and potential roles in host-microbe interactions. Here, we demonstrate how inverse stable isotope labeling (InverSIL) can rapidly link predicted siderophore gene clusters to their corresponding metabolites. By applying InverSIL to diverse bacterial strains, we validate known siderophore products and uncover unexpected products, highlighting the limitations of current *in silico* predictions. This study highlights the value of combining experimental approaches with genome mining to advance our understanding of how bacteria interact with each other and their environment.

## INTRODUCTION

Iron is an essential micronutrient for nearly all bacteria, yet it can be difficult to access for both free-living and host-associated species (1–3). To overcome this limitation, bacteria produce siderophores, secondary metabolites that chelate Fe(III) and facilitate its uptake (4, 5). Siderophores are typically constructed with a conserved biosynthetic logic via two major pathways, nonribosomal peptide synthetase (NRPS)-dependent and NRPS-independent siderophore (NIS) pathways. NRPS-dependent siderophores are built by large, modular enzyme complexes that assemble complex structures through condensation of amino acid building blocks (4). In contrast, NIS siderophores are assembled by smaller enzymes, mainly IucA- and IucC-like proteins, which perform the activation and condensation of different building blocks (6).

In bacteria, the genes responsible for the biosynthesis and transport of secondary metabolites, including siderophores, are typically colocalized and organized into biosynthetic gene clusters (BGCs) (7, 8). Advances in genome sequencing and computational tools such as antiSMASH (9) and PRISM (10) have made it increasingly straightforward to predict BGCs in bacterial genomes. However, bioinformatics predictions alone are often insufficient for identifying the specific chemical structures of BGC products due to factors such as enzyme promiscuity and crosstalk among different gene clusters (11–13). Strategies have also been developed for identifying siderophores by detecting iron-bound complexes by mass spectrometry (14, 15), however it can be difficult to rapidly link metabolomic “dark matter” to a specific BGC of interest (16). Most siderophore BGCs therefore remain unlinked to their corresponding products, which limits our understanding of bacterial iron acquisition strategies.

Strategies for integrating metabolomic and genomic datasets have been developed to improve researchers’ ability to identify the products of BGCs. For example, researchers can use stable isotope labeling to trace precursor incorporation into secondary metabolites, enabling linkage of BGCs to their products in a gene-to-molecule approach. This strategy, sometimes referred to as the genomisotopic approach, uses pathway-specific isotopically substituted precursors predicted from BGCs (17, 18). However, one limitation of isotopic labeling strategies is the cost and availability of isotopically substituted precursors.

Recently, we demonstrated that inverse stable isotope labeling (InverSIL) is a useful strategy for linking BGCs to their corresponding products (19, 20). This approach utilizes precursors at their natural isotopic abundance (referred to here as ^12^C for simplicity) in a ^13^C-substituted carbon background to detect precursor incorporation (21, 22). For example, a bacterial culture can be grown with ^13^C-subtituted glucose [(^13^C)glucose] as the sole carbon source, and then fed ^12^C-precursors so that their incorporation can be detected by mass spectrometry. This approach offers greater versatility than traditional isotopic labeling methods that rely on the availability of ^13^C-substituted precursors.

Catecholate siderophores have some of the highest affinities for Fe(III), by coordinating the iron atom through bidentate interactions between catechol hydroxylate groups in dihydroxybenzoic acid (DHB) (5, 23, 24). Two regioisomers of DHB, 2,3-dihydroxybenzoic acid (2,3-DHB) and 3,4-dihydroxybenzoic acid (3,4-DHB), are commonly incorporated into catecholate siderophores. Both are derived from the shikimate pathway, a central metabolic pathway connecting primary and secondary metabolism in bacteria. Specifically, 2,3-DHB is synthesized from chorismate via the enzymes encoded in the *dhb* operon (25), while 3,4-DHB is produced from 3-dehydroshikimate by the dehydratase AsbF (26, 27) (**Figure 1A**). Given the conserved roles of these enzymes in catecholate siderophore biosynthesis, we can use their corresponding genes to identify candidate siderophore BGCs from underexplored microbes. Inverse labeling through incorporation of DHB provides a predictable 7-Da isotopic downshift, which is readily detectable by automated mass spectrometry workflows (28) (**Figure 1B**). Furthermore, to our knowledge, 2,3-DHB is not currently commercially available in an isotopically substituted form, highlighting the utility of this approach for siderophore discovery. Using this method, we recently identified a new triscatecholate siderophore called methylocystabactin that is produced by many methane-oxidizing alphaproteobacteria (29).

**Figure 1.**
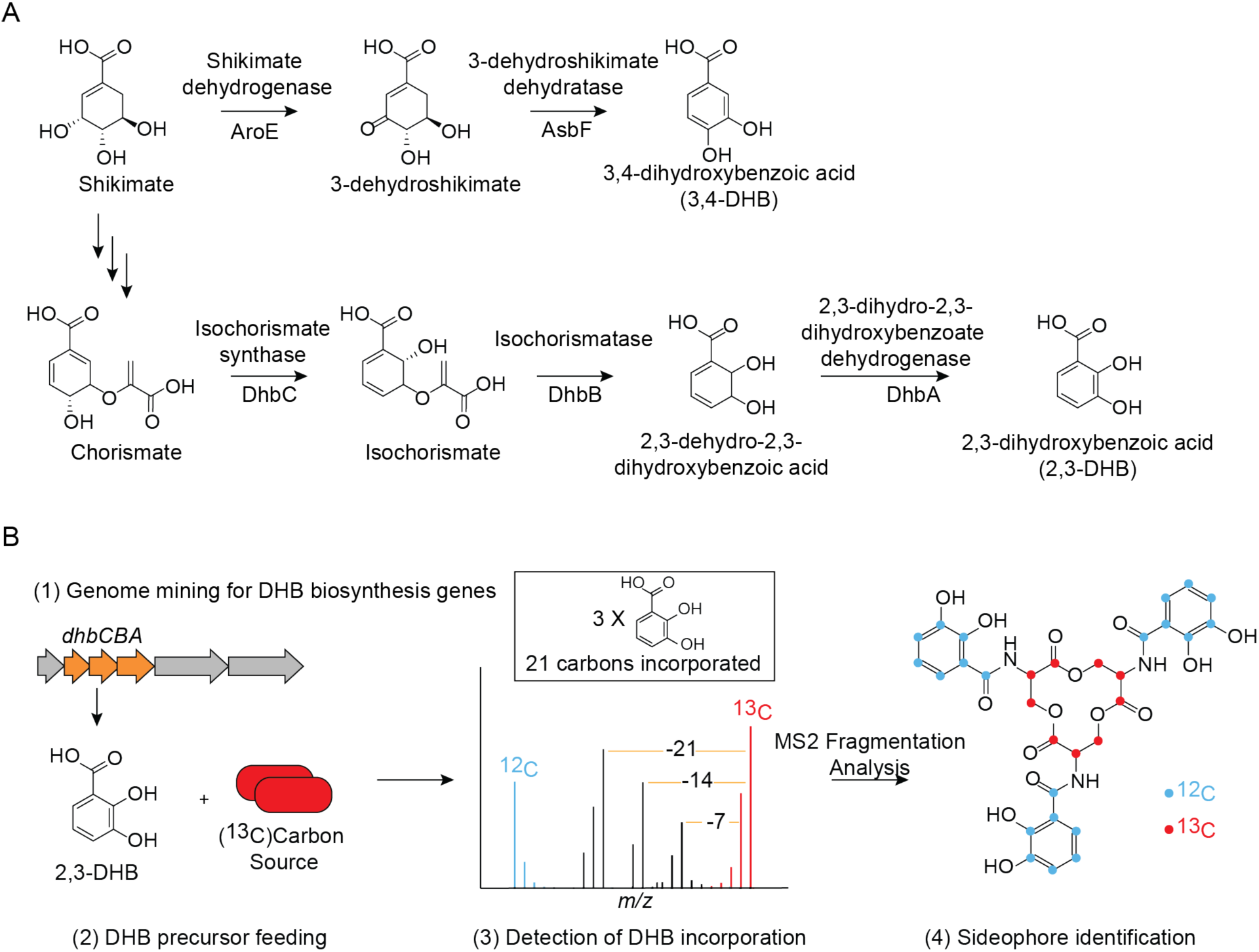
Inverse stable isotope labeling (InverSIL) for linking predicted catecholate siderophore BGCs with their products. (A) Enzymes responsible for the biosynthesis of catechols 3,4-DHB and 2,3-DHB. (B) Scheme of an InverSIL experiment: (1) Genome mining for DHB biosynthesis genes; (2) Addition of 2,3-DHB at its natural isotopic abundance to a bacterial culture grown on a ^13^C-substituted carbon source [indicated as (^13^C)]; (3) High-resolution mass spectrometry to detect DHB incorporation; (4) Identification of the siderophore aided by MS2 fragmentation. The siderophore enterobactin is shown as an example.

In this study, we applied InverSIL as a gene-to-molecule strategy to identify and structurally characterize the catecholate siderophore products of predicted BGCs in diverse free-living and host-associated bacterial genera. We used the precursors 2,3-DHB and 3,4-DHB to link predicted siderophore BGCs to their products in the methylotrophic genera *Methylophilus* and *Methylobacterium*, as well as to determine the structures of two siderophores previously shown to be essential for virulence by the opportunistic pathogen *Chromobacterium violaceum*. Next, we used InverSIL to identify the production of the siderophore enterobactin in two genera, *Kushneria* and *Paracoccus*, which was difficult to predict from genomic information due to the distributed nature of the siderophore biosynthetic genes. Finally, we used InverSIL to characterize new siderophores, cellulochelin A and B, from the cellulolytic plant symbiont *Cellulomonas* sp. strain Leaf334. Collectively, these findings demonstrate the value of experimental approaches like InverSIL for uncovering cryptic biosynthetic logic and expanding our understanding of iron acquisition in bacteria.

## RESULTS

### InverSIL links both predicted NRPS and NIS BGCs with their siderophore products

We first wanted to determine if the InverSIL approach could be applied to both predicted NRPS-dependent and NIS siderophore BGCs. By searching for the *dhb* genes in bacterial genomes, we identified a predicted NRPS-dependent siderophore BGC that was conserved within the genomes of several strains of the *Methylophilus* genus of methylotrophic bacteria. *Methylophilus* are betaproteobacteria that play an important role in the carbon cycle (30, 31), yet to our knowledge how they obtain iron from the environment has remained undetermined.

The predicted NRPS BGC has high similarity with the BGC that encodes for the production of the biscatecholate siderophore cepaciachelin (32), also known as protochelin C (33) **(Figure 2A)**, leading us to hypothesize that the *Methylophilus* BGC may produce the same compound. To test this hypothesis, we performed an InverSIL experiment with *Methylophilus* sp. strain 5 (34). We grew this strain with Fe(III) as the sole iron source and (^13^C)methanol as the sole carbon source, adding the precursor 2,3-DHB at its natural isotopic abundance to one sample. InverSIL revealed a metabolite incorporating two 2,3-DHB units with the same high-resolution mass and carbon count as cepaciachelin **(Figures 2B, 2C)**. Tandem MS (MS2) fragmentation and direct comparison with extracts from the known cepaciachelin producer *Burkholderia ambifaria* BAA-244 (also known as strain AMMD) (32), confirmed that *Methylophilus* sp. strain 5 produces cepaciachelin (**Figures 2C, 2D**, **Table S1**). In addition to cepaciachelin, we detected azotochelin and aminochelin (**Figure S1**), two structurally related compounds that have also previously been described as biosynthetic intermediates in protochelin biosynthesis (33).

**Figure 2.**
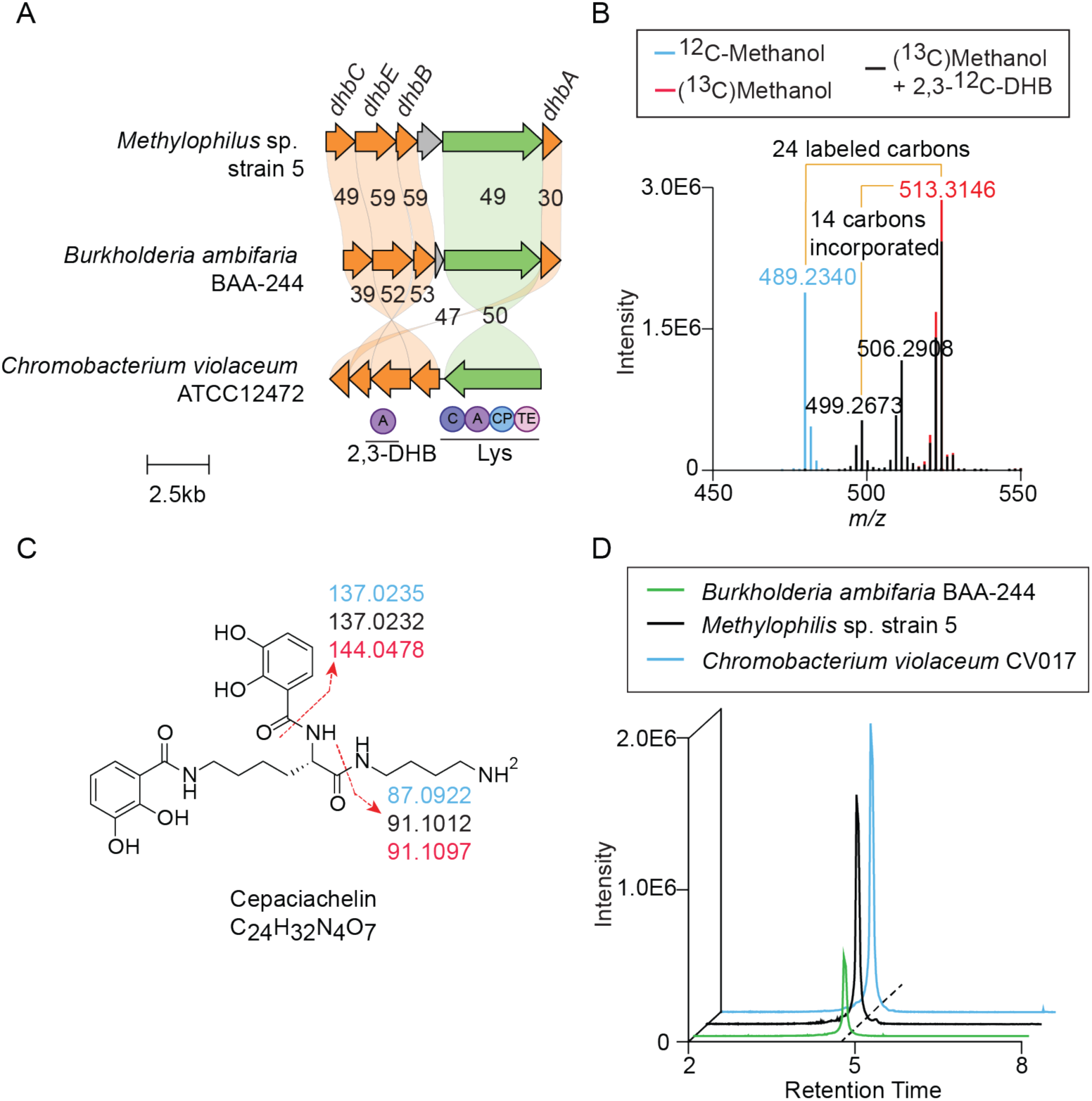
Using InverSIL to link an NRPS-dependent catecholate siderophore BGC with its product, cepaciachelin. (A) Comparison of an NRPS-dependent BGC in *Methylophilus* sp. strain 5, *Burkholderia ambifaria* BAA-244, and *Chromobacterium violaceum* CV017. Arrow colors indicate predicted 2,3-DHB biosynthesis genes (orange), predicted NRPS genes (green), and other genes (gray). Circles indicate predicted NRPS domains C (condensation domain), A (adenylation domain), CP (acyl-carrier protein), and TE (thioesterase domain), with the predicted adenylation domain substrates listed below. (B) Overlayed mass spectra of *Methylophilus* sp. strain 5 supernatant extract showing incorporation of two 2,3-DHB units into a metabolite with the same high-resolution mass and carbon count as cepaciachelin. (C) Structure of cepaciachelin showing MS2 fragments from different InverSIL conditions. The colors match the growth conditions indicated in panel B. (D) Extracted ion chromatograms of supernatant extracts of *B. ambifaria* BAA-244, *Methylophilus* sp. strain 5, and *Chromobacterium violaceum* CV017 for *m/z* 489.2340, corresponding to the [M+H]^+^ of cepaciachelin. Mass tolerance < 5ppm.

To further test the capability of InverSIL to link catecholate siderophore BGCs to their products, we targeted a NIS BGC that is widespread in pink pigmented facultative methylotrophs of the genera *Methyobacterium* and *Methylorubrum*. These methylotrophs often promote plant growth (35) and can cause hospital-acquired infections in some instances (36). The NIS BGC contains an *asbF* homolog and is similar to the BGCs for the production of the recently described lanthanide-chelating molecule methylolanthanin (37), as well as rhodopetrobactin (38), a siderophore known to utilize 3,4-DHB as its iron-chelating moiety (**Figure 3A**).

**Figure 3.**
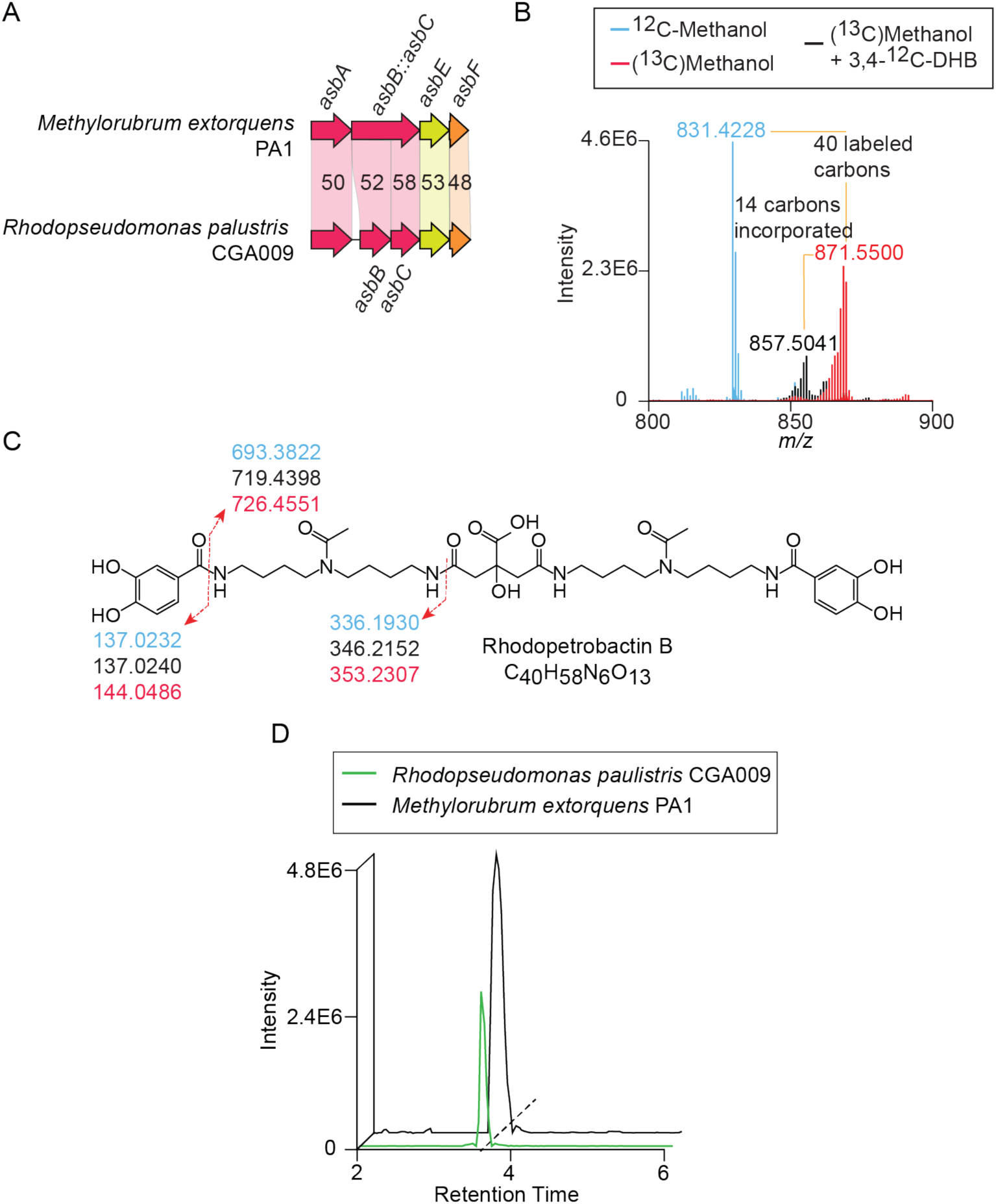
Using InverSIL to link a NIS catecholate siderophore BGC with its product, rhodopetrobactin B. (A) Comparison of a NIS siderophore BGC in *Methylorubrum extorquens* PA1 and *Rhodopseudomonas palustris* CGA009. Arrow colors indicate predicted *iucA*/*iucC* genes (red), a conserved hypothetical protein in petrobactin biosynthesis (yellow), and a predicted *asbF* gene (orange). (B) Overlayed mass spectra of *M. extorquens* PA1 supernatant extract showing incorporation of two 3,4-DHB units into a metabolite with the same high-resolution mass and carbon count as rhodopetrobactin B. (C) Structure of rhodopetrobactin B showing MS2 fragments from different InverSIL conditions. The colors match the growth conditions indicated in panel B. (D) Extracted ion chromatograms of supernatant extracts of *R. palustris* CGA009 and *M. extorquens* PA1 for *m/z* 831.4228, corresponding to the [M+H]^+^ of rhodopetrobactin B. Mass tolerance < 5ppm.

To determine the product of this BGC, we grew *Methylorubrum extorquens* PA1 on (^13^C)methanol, and added 3,4-DHB at its natural isotopic abundance to one sample. InverSIL revealed a compound incorporating two 3,4-DHB units with a high-resolution mass and carbon count identical to that of rhodopetrobactin B (**Figure 3B**). We further confirmed this with MS2 fragmentation and direct comparison with extracts from the known rhodopetrobactin producer *Rhodopseudomonas palustris* CGA009 (38) (**Figures 3C, 3D**, **Table S2**). We did not detect methylolanthanin production by *M. extorquens* PA1 (data not shown). Together, these results demonstrate the utility of InverSIL for determining the siderophore products of both predicted NRPS and NIS BGCs.

### InverSIL identifies the structures of two siderophores produced by the opportunistic pathogen *Chromobacterium violaceum*

Next, we examined the genome of the opportunistic pathogen *C. violaceum,* which has been reported to produce two catecholate siderophores essential for virulence that were named chromobactin and viobactin (39). To date, these siderophores have only been characterized genetically and through phenotypic assays with *C. violaceum* ATCC12472 (39), providing an opportunity to determine the siderophore structures directly using InverSIL. The chromobactin BGC contains the *dhb* genes for 2,3-DHB biosynthesis and is homologous to the cepaciachelin BGC in *Methylophilus* sp. strain 5 **(Figures 2A, 4A)**. The viobactin BGC resembles a triscatecholate siderophore BGC predicted to incorporate a cationic amino acid spacer (40), as seen in the siderophore cyclic trichrysobactin (41) (**Figure 4A**).

**Figure 4.**
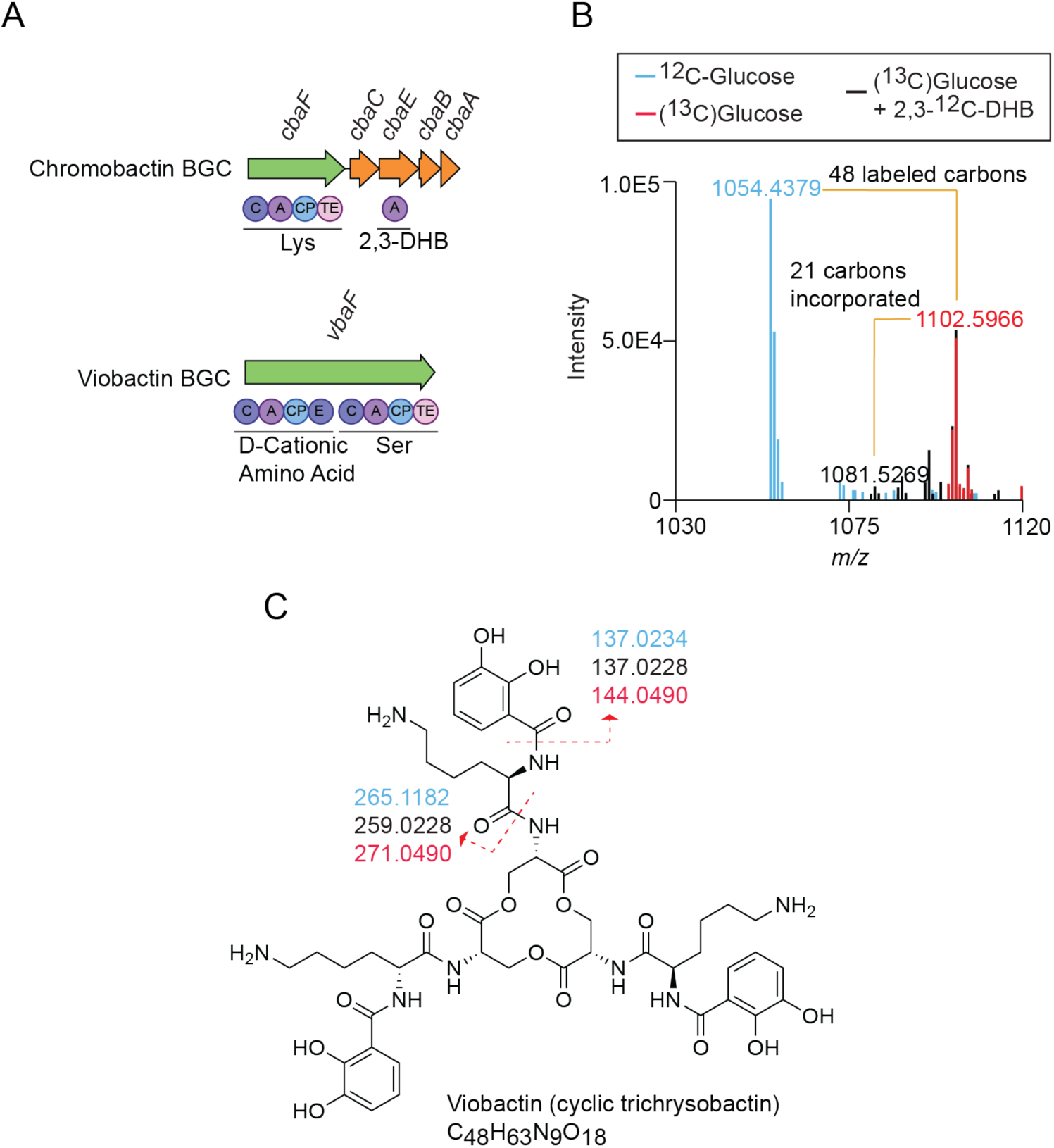
Using InverSIL to determine the structures of the siderophores used by the opportunistic pathogen *Chromobacterium violaceum*. (A) BGCs for the *C. violaceum* siderophores. Arrow colors indicate predicted NRPS genes (green) and 2,3-DHB biosynthesis genes (orange). Circles indicate predicted NRPS domains C (condensation domain), A (adenylation domain), CP (acyl-carrier protein), TE (thioesterase domain), and E (epimerization domain), with the predicted adenylation domain substrates listed below. (B) Overlayed mass spectra of *C. violaceum* CV017 supernatant extract showing incorporation of three 2,3-DHB units into a metabolite with the same high-resolution mass and carbon count as cyclic trichrysobactin. (C) Structure of viobactin (cyclic trichrysobactin) showing MS2 fragments from different InverSIL conditions. The colors match the growth conditions indicated in panel B.

To determine the structures of chromobactin and viobactin, we performed InverSIL on *C. violaceum* CV017, which contains both BGCs, using 2,3-DHB at its natural isotopic abundance as the precursor in M9 minimal medium with (^13^C)glucose as the sole carbon source. Consistent with the homology of the chromobactin BGC with the cepaciachelin BGC in *Methylophilus* sp. strain 5, we identified a metabolite with the same high-resolution mass and carbon count as cepaciachelin that incorporated two 2,3-DHB units **(Figure S2).** We confirmed the identity of cepaciachelin using MS2 fragmentation and comparison with extracts from *Methylophilus* sp. strain 5 and *B. ambifaria* BAA-244 (**Figures 2C, 2D, 4B**; **Table S1**). The siderophore called chromobactin is therefore structurally identical to cepaciachelin.

For the product of the viobactin BGC, we detected a metabolite that incorporated three 2,3-DHB units, consistent with a triscatecholate siderophore **(Figure 4C)**. The high-resolution mass, carbon count, MS2 fragmentation, and Marfey’s analysis identified this compound as cyclic trichrysobactin, which contains a D-lysine cationic amino acid spacer, as predicted from the epimerization domain in the NRPS gene of the viobactin BGC (**Figures 4C, 4D**, **S3, S4**, **Tables S3, S4**) (41). This confirms that the viobactin BGC produces a siderophore structurally identical to cyclic trichrysobactin. These results highlight the utility of InverSIL for rapid structural characterization of siderophores with biomedical relevance.

### InverSIL uncovers enterobactin production by the genera *Kushneria* and *Paracoccus* despite distributed biosynthetic genes

Members of the *Kushneria* genus are plant endophytes and have been explored for potential to promote plant growth (42, 43), however, how these bacteria acquire iron has remained unknown to our knowledge. We identified a BGC primarily associated with *Kushneria* species, where the *dhb* biosynthetic operon is colocalized with a NIS gene cluster with similarity to the aerobactin BGC (**Figure 5A**). Based on these features, we hypothesized that this BGC may encode for the production of a novel NIS siderophore with 2,3-DHB chelating groups instead of the typical 3,4-DHB moieties.

**Figure 5.**
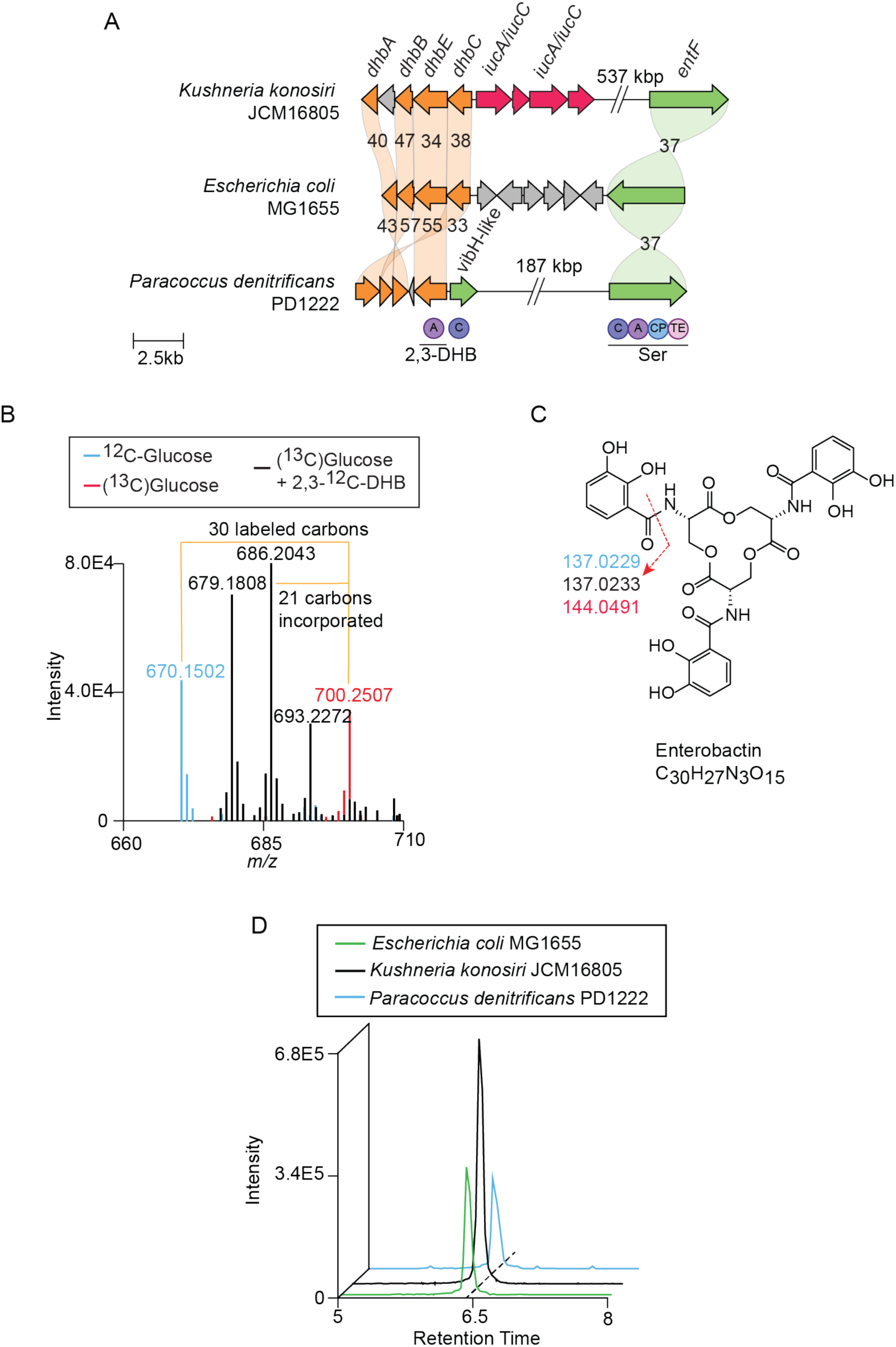
Non-colocalized enterobactin BGCs in *Kushneria konosiri* JCM16805 and *Paracoccus denitrificans* PD1222. (A) Comparison of a predicted *dhb* operon, NIS BGC, and distant *entF* gene in *K. konosiri* JCM16805 with the enterobactin BGC in *E. coli* MG1655 and another non-colocalized enterobactin BGC in *Paracoccus denitrificans* PD1222. Arrow colors indicate predicted 2,3-DHB biosynthesis genes (orange), predicted *iucA*/*iucC* genes (red), predicted NRPS genes (green), and other genes (gray). Circles indicate predicted NRPS domains C (condensation domain), A (adenylation domain), CP (acyl-carrier protein), and TE (thioesterase domain), with the predicted adenylation domain substrates listed below. (B) Overlayed mass spectra of *K. konosiri* JCM16805 supernatant extract showing incorporation of three 2,3-DHB units into a metabolite with the same high-resolution mass and carbon count as enterobactin. (C) Structure of enterobactin showing MS2 fragments from different InverSIL conditions. The colors match the growth conditions indicated in panel B. (D) Extracted ion chromatograms of supernatant extracts from *K. konosiri* JCM 16805, *P. denitrificans* PD1222, and *E. coli* MG1655 for *m/z* 670.1502, corresponding to the [M+H]^+^ of enterobactin. Mass tolerance < 5ppm.

To test this hypothesis, we selected *Kushneria konosiri* JCM16805 and performed InverSIL using 2,3-DHB at its natural isotopic abundance as the precursor in cultures grown on M9 minimal medium with (^13^C)glucose as the sole carbon source. Inverse labeling revealed a compound incorporating three 2,3-DHB units (**Figure 5B**). The high-resolution mass, carbon count, and MS2 fragmentation indicated that this compound is the well-characterized triscatecholate siderophore enterobactin (**Figures 5C**). Direct comparison to an extract from *Escherichia coli* MG1655 confirmed that *K. konosiri* JCM16805 indeed produces enterobactin (**Figure 5D**, **Table S5**). We also detected linear dimer and trimer forms of enterobactin, both of which were supported by precursor incorporation patterns and MS2 fragmentation (**Figure S5**). Additionally, we detected aerobactin, which is the predicted product of the NIS BGC (**Figure S6, Table S6**). This result was unexpected because genomic predictions did not indicate the presence of an enterobactin biosynthetic pathway in this strain. Further analysis of the *K. konosiri* JCM16805 genome revealed that an *entF* homolog, essential for enterobactin biosynthesis, is located in a different part of the genome, more than 500 kilobase pairs from the *dhb* operon (**Figure 5A**).

Based on this finding, we discovered another *entF* homolog in the genome of *Paracoccus denitrificans* PD1222 that is distantly located from the *dhb* genes for 2,3-DHB biosynthesis (**Figure 5A**). *P. denitrificans* PD1222 has been well studied for its role in the nitrogen cycle and is known to produce the siderophore parabactin, a mixed-ligand siderophore containing two catecholates from 2,3-DHB and one phenolate from 2-hydroxybenzoic acid (salicylic acid) (44–46). *P. denitrificans* PD1222 also produces the siderophore brucebactin, which is considered an intermediate in parabactin biosynthesis (45, 47). However, to our knowledge production of enterobactin by *P. denitrificans* PD1222 has not been previously reported.

To determine if *P. denitrificans* PD1222 could also produce enterobactin, we grew the strain in M9 minimal medium and performed InverSIL using 2,3-DHB and salicylic acid as precursors and (^13^C)glucose as the sole carbon source. As expected, we detected both parabactin and brucebactin based on high-resolution mass, carbon count, precursor incorporation and MS2 fragmentation (**Figures S7-S8**). Notably, we also detected incorporation of 2,3-DHB into a triscatecholate siderophore we identified as enterobactin (**Figure S9**). To our knowledge, this is the first report of enterobactin production by *P. denitrificans* PD1222. The fragmented and unannotated gene arrangement highlights the utility of experimental approaches to reveal functionally complete siderophore pathways distributed across bacterial genomes.

### InverSIL identifies cellulochelin, a new siderophore produced by *Cellulomonas* sp. strain Leaf334

Finally, we used InverSIL to characterize a new siderophore. We applied the same genome-mining strategy used in earlier experiments, focusing on the presence of the *dhb* genes as indicators of catecholate siderophore biosynthesis. This search identified a predicted NRPS-dependent siderophore BGC in *Cellulomonas* sp. strain Leaf334, a cellulolytic actinobacterium isolated from the model plant *Arabidopsis thaliana* (48) (**Figure 6A**).

**Figure 6.**
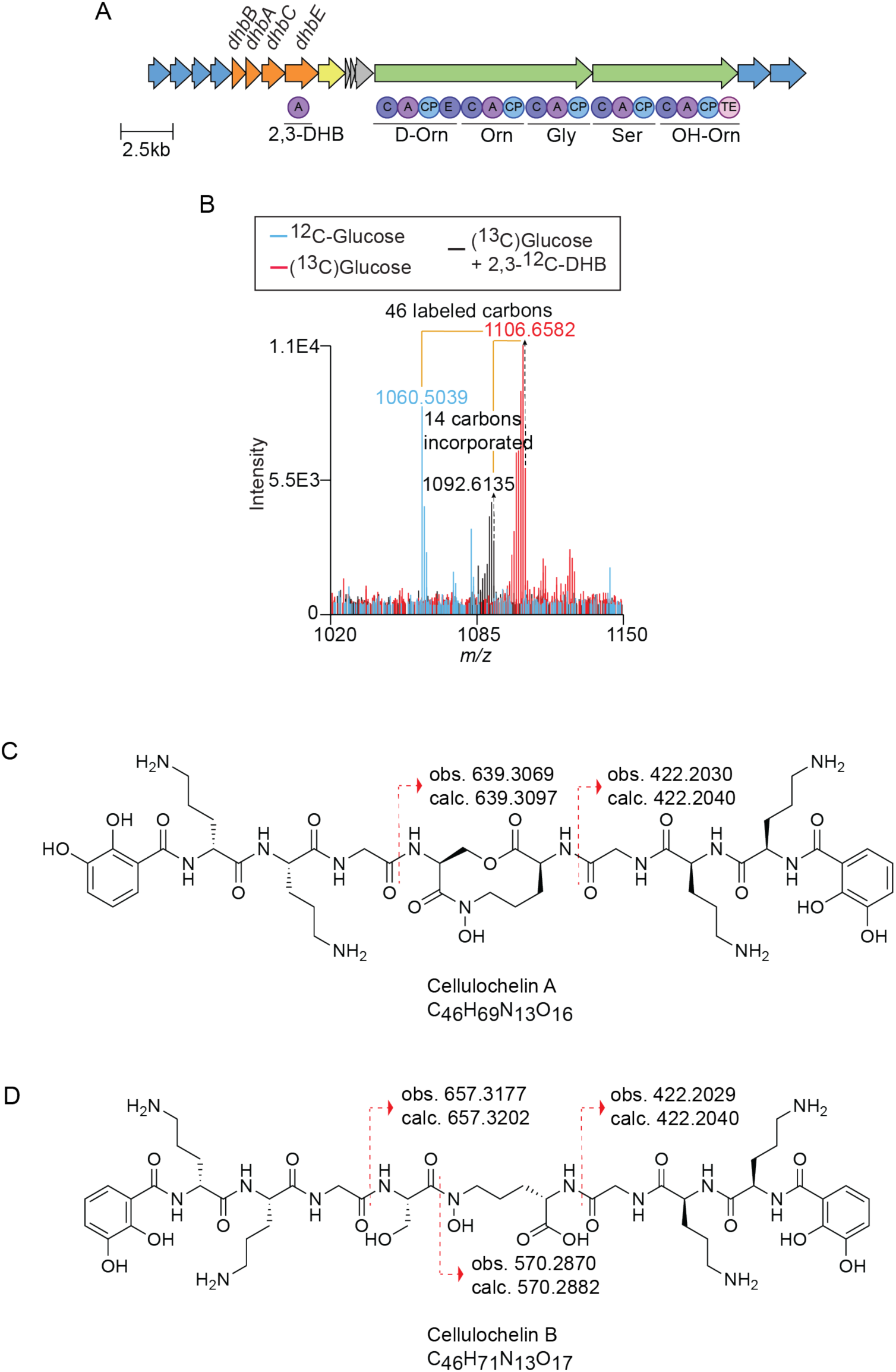
Characterization of a novel siderophore from *Cellulomonas* sp. strain Leaf334 using InverSIL. (A) The cellulochelin BGC. Arrow colors indicate predicted 2,3-DHB biosynthesis genes (orange), predicted NRPS genes (green), predicted transport-related genes (blue), and a predicted ferric reductase gene (yellow). Circles indicate predicted NRPS domains C (condensation domain), A (adenylation domain), CP (acyl-carrier protein), and TE (thioesterase domain), and E (epimerization domain), with the predicted adenylation domain substrates listed below. (B) Overlayed mass spectra of *Cellulomonas* sp. strain Leaf334 supernatant extract showing incorporation of two 2,3-DHB units into a new metabolite. (C,D) Structure of (C) cellulochelin A and (D) cellulochelin B showing diagnostic MS2 fragments for differentiating the two siderophores.

Genome analysis revealed that this BGC encodes the *dhb* operon including *dhbE* and two predicted NRPS genes (**Figure 6A**). The first NRPS gene contains three modules predicted to incorporate D-ornithine, L-ornithine, and glycine. The second NRPS gene contains two modules followed by a thioesterase domain and is predicted to add L-serine and *N*-hydroxy-L-ornithine. This second NRPS gene shares homology with both *fscH* and *fscI* from *Thermobifida fusca*, which together form the 10-membered cyclic hydroxamate seryl-ester in the siderophore fuscachelin (49). Fuscachelin has a unique heterodimeric architecture assembled through a non-linear NRPS pathway, and contains two terminal 2,3-DHB units and a hydroxamate moiety, with a heterodimeric peptide backbone composed of arginine and two glycines (49).

Based on the similarity of the BGC architecture and domain predictions, we hypothesized that *Cellulomonas* sp. strain Leaf334 produces a siderophore featuring two terminal DHB units and a 10-membered cyclic hydroxamate ester derived from serine and *N*-hydroxy-L-ornithine. To test this hypothesis using InverSIL, we grew *Cellulomonas* sp. strain Leaf334 in M9 minimal medium with (^13^C)glucose as the sole carbon source, and 2,3-DHB at its natural isotopic abundance as the precursor. This revealed a metabolite incorporating two 2,3-DHB units (**Figure 6B**), which, to our knowledge, does not match the high-resolution mass of any known siderophore. Targeted MS2 fragmentation of the detected metabolite indicated it is a peptide containing DHB-Orn-Orn-Gly-Ser-OHOrn-Gly-Orn-Orn-DHB, which matched with the predicted substrates of the NRPS gene (**Figures 5C, S10, S11**). We named this new siderophore cellulochelin A.

To confirm the structure of cellulochelin A, we scaled up cultures of *Cellulomonas* sp. Leaf334 to purify sufficient material for NMR analysis. Like fuscachelin A, cellulochelin A contains a 10-membered macrolactone ring that undergoes dehydration to form cellulochelin B (**Figure 5D, S11**). This conversion is supported by the presence of a serine and *N*-hydroxy-ornithine MS2 fragment in cellulochelin B, which are absent in cellulochelin A (**Figure 5D, S10, S11, Table S9, S10**). We performed NMR and Marfey’s analysis on cellulochelin B, confirming the overall structure and the presence of the predicted D-ornithine in the structure (**Figure S12-S17**; **Table S7, S8**). We were unable to isolate enough cellulochelin A for full NMR characterization because it readily converts to cellulochelin B, as previously reported for fuscachelin A (49). Comparison of MS2 fragmentation data for cellulochelin A and B supports the predicted relationship between these two compounds, showing an MS2 neutral loss consistent with the presence of a macrolactone ring in cellulochelin A, which is absent in cellulochelin B (**Figures 5C, 5D, S10, S11**). The cellulochelins represent a new addition to the fuscachelin-like siderophores, expanding the structural diversity of microbial iron-chelating compounds.

## DISCUSSION

Understanding which siderophores bacteria produce is critical for elucidating microbial iron acquisition strategies and ecological interactions with other strains and/or the host (2, 3). Here, we used InverSIL to rapidly link predicted siderophore BGCs with their catecholate siderophore products in diverse free-living and host-associated bacteria. We show this approach is effective across multiple siderophore biosynthetic pathways, including NRPS-dependent and NIS systems, and can help elucidate complex biosynthetic pathways involving crosstalk between different clusters.

Our findings underscore the importance of experimentally validating bioinformatic predictions and highlight unanswered questions about metallophore biosynthesis. For example, the NIS BGC in *M. extorquens* PA1 produced the siderophore rhodopetrobactin B but not the lanthanide-chelating compound methylolanthanin, although this BGC is highly similar to the one in *Methylorubrum extorquens* AM1 that produces methylolanthanin (37). Further work will be needed to resolve this discrepancy and potential biosynthetic divergence between methylolanthanin and rhodopetrobactin production in pink pigmented facultative methylotrophs.

InverSIL also revealed unexpected siderophore production from fragmented BGCs in bacterial genomes. In *K. konosiri* JCM16805 and *P. denitrificans* PD1222, enterobactin was detected despite the absence of a canonical enterobactin BGC. In both cases, we identified a distantly located *entF* homolog, highlighting that functional siderophore pathways can persist even when distributed across the genome. Enterobactin production and presumed utilization is notable because this siderophore is unparalleled in its affinity for iron (24) and can be produced by diverse species found in soils (50). Multiple siderophore production is prevalent across diverse bacterial taxa and often involves sharing of precursors such as 2,3- and 3,4-DHB between distinct biosynthetic pathways (12, 51). The production of diverse siderophores by one bacterial strain may be due to myriad reasons including optimized uptake of different metals found at different environmental concentrations, and/or microbial competition (52).

We also used InverSIL to determine the identities of two siderophores produced by the opportunistic pathogen *C. violaceum* that are known to be important for virulence (39). We determined that the products of these BGCs, previously named chromobactin and viobactin, are identical to cepaciachelin and cyclic trichrysobactin, respectively. It is interesting that these same siderophores are produced by other pathogens including *Burkholderia* spp. of human pathogens for cepaciachelin (32) and *Dickeya* spp. of plant pathogens for cyclic trichrysobactin (41). Iron availability is a critical factor in microbial pathogenesis (2, 3), and knowing the molecular details of iron acquisition by *C. violaceum* may provide new insight into infections by this species.

Finally, we used InverSIL to characterize cellulochelin A and cellulochelin B, new siderophores from *Cellulomonas* sp. strain Leaf334. Cellulochelin A features a 10-membered cyclic hydroxamate ester, similar to that found in fuscachelin A. The discovery of cellulochelin expands the chemical diversity of microbial iron-chelating compounds. Further studies will be needed to determine the ecological role of the cellulochelins, especially in the context of its association with its host *Arabidopsis thaliana*.

In summary, InverSIL provides a powerful and versatile platform for experimentally linking siderophore BGCs to their products, thereby complementing genome-based predictions. By enabling rapid structural elucidation, this approach not only expands our understanding of siderophore diversity and biosynthetic logic but also accelerates the discovery of secondary metabolites with ecological and biotechnological significance. The InverSIL approach can also be extended to identify secondary metabolites that contain other functional groups of interest, enabling the linkage of more bacterial biosynthetic genes to their products.

## ACKNOWLEDGEMENTS

This work was supported by National Institutes of Health grant R35 GM147018 (to A.W.P.). T.C.E.L. was supported by NIH training grant T32 AI055434. We thank C.S. Harwood and A.L. Schaefer (University of Washington) for providing *R. palustris* CGA009 and *C. violaceum* CV017, and T.L. Karasov (University of Utah) for providing *Cellulomonas* sp. strain Leaf334. We thank A.T. Aron (University of Denver) and members of the Puri Lab for helpful discussions.

## DATA AVAILABILITY

The mass spectrometry data were deposited in the public repository MassIVE via ID MSV000099756.

## AUTHOR CONTRIBUTIONS

J.M.D.R. and A.W.P. designed the experiments. J.M.D.R., T.C.E.L., and V.P.M. performed the experiments. J.M.D.R., T.C.E.L., and A.W.P. analyzed the data. J.M.D.R. and A.W.P. wrote the manuscript. All authors edited and approved of the final version of the manuscript.

## METHODS

### General Experimental Procedures

NMR spectra were obtained in D_2_O (δH 4.66 ppm) using an Agilent DirectDrive 500 with a high-sensitivity cold probe detection system. Reverse-phase HPLC was performed using an Agilent 1260 Infinity HPLC system with a Waters Sunfire C_18_ OBD =column (5 μm, 10 × 100 mm). Isotopically substituted compounds were purchased from Cambridge Isotope Laboratories.

### Routine Bacterial Culturing

All strains used in the study is summarized in **Table S11**. *Kushneria konosiri* JCM 16805 was grown at 30°C in the medium described by Ventosa et al. (53). This medium contains 75 g L^-1^ NaCl, 2 g L^-1^ KCl, 0.2 g L^-1^ MgSO_4_. • 7H_2_O, 1 g L^-1^ KNO_3_, 1 g L^-1^ (NH_4_)_2_HPO_4_, 0.5 g L^-1^ KH_2_PO_4_, 0.05 g L^-1^ yeast extract, and 2 g L^-1^ ^12^C- or ^13^C-glucose. *Escherichia coli* MG1655, *Burkholderia ambifaria* BA-244, *Chromobacter violaceum* CV017, *Cellulomonas* sp. strain Leaf334, and *Paracoccus denitrificans* PD1222 were grown in M9 minimal medium supplemented with 4.0 g L^-1^ glucose or ^13^C-glucose. M9 minimal medium consists of 12.8 g L^-1^ Na_2_HPO_4_ • 7H_2_O, 3 g L^-1^ KH_2_PO_4_, 0.5 g L^-1^ NaCl, 1.0 g L^-1^ NH_4_Cl, 0.2408 g L^-1^ MgSO_4_, and 0.011 g L^-1^ CaCl_2_. *E. coli* MG1655 was grown at 37°C. *C. violaceum* CV017, *B. ambifaria* BAA-244, *P. denitrificans* PD1222 and *Cellulomonas* sp. strain Leaf334 were grown at 30°C. For *Cellulomonas* sp. strain Leaf334, a final concentration of 1% (v/v) yeast extract was added to the M9 medium. *Methylorubrum extorquens* PA1 and *Methylophilus* sp. strain 5 were grown in nitrate mineral salts (NMS) medium. NMS medium contains 0.2 g/l MgSO4·7H2O, 0.2 g/l CaCl_2_·6H2O, 1 g/l KNO_3_, 30 μM LaCl3, a final concentration of 5.8-mM phosphate buffer (pH 6.8), and 1X trace elements; 500X trace elements contains 1.0 g/l Na2-EDTA, 2.0 g/l FeSO4·7H2O, 0.8 g/l ZnSO4·7H2O, 0.03 g/l MnCl2·4H2O, 0.03 g/l H3BO3, 0.2 g/l CoCl2·6H2O, 0.6 g/l CuCl2·2H2O, 0.02 g/l NiCl2·6H2O, and 0.05 g/l Na2MoO·2H2O. *Rhodopseudomonas palustris* CGA009 was grown in photosynthetic medium as previously described (54). All cultures were shaken at 200 rpm, 30°C.

### High Resolution Tandem Mass Spectrometry (LC-HRMS/MS)

Mass spectrometry data were collected using an Agilent Revident Q-TOF coupled to an Agilent Infinity III HPLC system with an Acquity UPLC HSS T3 C18 column (1.8 μm, 2.1 x 50 mm). Solvent A: Water + 0.1 % (v/v) formic acid, Solvent B: Acetonitrile + 0.1% (v/v) formic acid. The sample was eluted from the column using a 12-minute linear solvent gradient: 0-0.1 min, 1% B; 0.1 - 10 min, 1-95% B. The solvent flow rate was 0.4 mL min^-1^. Mass spectra were collected in positive ion mode, using the Auto-MS^2^ acquisition mode. An MS range of *m/z* 100-1500 and an MS2 range of *m/z* 50-1500, both at 5 spectra/s was used using the medium (∼ *m/z* 4) isolation width. The collision energy gradient was set to automatic according to *m/z* values of precursor ions. Under the collision energy section, Formula was used with two lines, charge set to “All”, “Slope” is 2.6 and “Offset” is 14.75, and charge set to “All”, “Slope” set to 3.9 and “Offset” to 22.13. The maximum precursors per cycle was set to 5, with the Absolute Precursor Threshold set to 15000 (Relative Threshold 0.015%) and Active Exclusion enabled.

### Genome Mining

Genome mining was done using the Joint Genome Institute Integrated Microbial Genomes and Microbiomes (JGI-IMG/M) system (55) using the gene search by function IDs using EC number (*dhbA* = EC 1.3.1.28, *dhbB* = EC 3.3.2.1, *dhbC* = EC 5.4.4.2, *asbF* = EC 4.2.1.118).

Strain genomes were download in NCBI Refseq and was ran on AntiSMASH v7.0 to further examine BGCs.

### Gene Clusters

The gene cluster shown for *Methylophilus* sp. strain 5 is from IMG gene ID 2516196715 to 2516196720. The gene cluster for *B. ambifaria* BAA244 is from IMG gene ID 2632565022 to 2632565027. The gene clusters for *C. violaceum* CV017 is from IMG gene ID 2728635959 to 2728635963 and 2728634501. The gene cluster for *M. extorquens* PA1 is from IMG gene ID 641369550 to 641369553. The gene cluster for *R. palustris* CGA009 is from IMG gene ID 637475518 to 637475522. The gene cluster for *E. coli* MG1655 is from IMG gene ID 8011516428 to 8011516438. The gene cluster for *K. konosiri* JCM16805 is from IMG gene ID 2752508317 to 2752508321, the *entF* homolog is IMG gene ID is 2752507833. The gene cluster for *P. denitrificans* PD1222 is from IMG gene ID 639769712 to 639769716, the *entF* homolog is IMG gene ID 639769893. The gene cluster for *Cellulomonas* sp. strain Leaf334 is from IMG gene ID 2645451482 to 2645451497. All gene IDs refer to the Joint Genome Institute the Joint Genome Institute Integrated Microbial Genomes and Microbiomes (JGI-IMG/M) system (55). BGCs were compared and visualized using CAGECAT (56).

### Inverse Stable Isotope Labeling

Inverse labeling experiments were performed as previously described (17). Briefly, exponentially growing cultures were first pelleted and resuspended in media with no carbon source. Subsequently, 2 mL cultures were inoculated into a final OD of 0.05, and the ^12^C-carbon source (methanol or glucose) was added to one culture, the ^13^C-substituted carbon source to another, and ^13^C-substituted carbon source plus 100 µM ^12^C-precursor (2,3-DHB or 3,4-DHB) to a final condition for the labeling experiment. All cultures were grown for five days, lyophilized, and resuspended in 200 µL 50% MeOH/H2O before being analyzed by LC-HRMS/MS. Datasets were subsequently analyzed as previously described (28).

### Cellulochelin B Purification and Structure Elucidation

To isolate sufficient quantities of cellulochelin B for structural elucidation, we grew large scale cultures of *Cellulomonas* sp. strain Leaf334. The presence of cellulochelin B was monitored throughout the purification by mass spectrometry. Briefly, an exponentially growing culture of *Cellulomonas* sp. strain Leaf334 was inoculated into 1000 mL of M9 minimal mediium supplemented with yeast extract and grown at 30°C and 200 rpm for 5 days for a total of 6.0 liters. After centrifugation, the cell-free supernatant was passed through a reverse phase C18 solid phase extraction (Waters Sep-pak C18, 5g) cartridge, activated with 100% methanol and conditioned with Milli-Q water. Cellulochelin B was eluted with 25% MeOH/H2O. Reverse-phase HPLC purification was performed using a Sunfire C18 OBD column (5 μm, 10 x 100 mm) using a gradient of 7%/93% to 20%/80% Methanol/H2O with 0.1% trifluoroacetic acid over 20 minutes at 4 mL/min while continuously monitoring the eluent at 310nm. Another reverse-phase HPLC using the same method was used to purify the compounds further (Cellulochelin B, 4.2 mg, t_R_ = 5.2 min).

### Cellulochelin B

C_46_H_71_N_13_O_17_. Light yellowish powder; High-Resolution MS: [M+H]+ calc. 1078.5164, obs. 1078.5150, −1.30 Δppm. NMR data: SI Appendix, **Figs. S12–S15** and **Table S7**.

### Viobactin (cyclic trichrysobactin) Purification

To confirm the stereochemistry of the lysine spacer in cyclic trichrysobactin, we grew *C. violaceum* CV017 in 100mL M9 minimal medium at 30°C and 200 rpm for 5 days. After centrifugation, cell-free supernatant was passed through a reverse phase C18 solid phase extraction (Waters Sep-pak C18, 1g) cartridge, activated with 100% methanol and conditioned with Milli-Q water. Chrysobactin was eluted with 50% MeOH/H2O. Reverse-phase HPLC purification was performed using a Sunfire C18 OBD column (5 μm, 10 x 100 mm) using a gradient of 5%/95% to 40%/60% Acetonitrile/H2O with 0.1% trifluoroacetic acid over 20 minutes at 4 mL/min while continuously monitoring the eluent at 310nm (Chrysobactin, 0.5mg t_R_ = 3 min). Fractions were injected to a high-resolution mass spectrometer to check for the presence of chrysobactin.

### Determination of Amino Acid Configuration

Purified cellulochelin B and cyclic trichrysobactin (∼1 mg) were hydrolyzed in 1 mL of 6 N HCl overnight at 110 °C with stirring. The solution was then evaporated to dryness, and the resulting solid was resuspended in 250 μL of water. An aliquot (50 μL) of the hydrolysate solution was then transferred to a clean glass vial to which 20 μL 1 M NaHCO_3_ and 50 μL Marfey’s reagent (L-FDLA, 1% w/v solution in acetone) were added. The mixture was stirred for 1 hour at 40°C and then quenched with 20 μL 1 N HCl. This solution was filtered and then dried before being resuspended in 50% MeOH/H2O for LC-HRMS analysis. The same derivatization using Marfey’s reagent was also performed on amino acid standards. Analysis was performed on Agilent Revident Q-TOF coupled to an Agilent Infinity III HPLC system with an Acquity UPLC HSS T3 C18 column (1.8 μm, 2.1 x 50 mm) using a gradient of 1%/99% ACN/H2O to 99%/1% ACN/H2O containing 0.1% formic acid, 30 minutes, 0.4 mL/min. Retention times for amino acids derived from cellulochelin, B, chrysobactin, and amino acids standards are summarized in **Tables S4, S8**.

## REFERENCES

1. Emerson D, Roden E, Twining B. 2012. The microbial ferrous wheel: iron cycling in terrestrial, freshwater, and marine environments. Front Microbiol 3.

2. Cassat JE, Skaar EP. 2013. Iron in Infection and Immunity. Cell Host & Microbe 13:509–519.

3. Kramer J, Özkaya Ö, Kümmerli R. 2020. Bacterial siderophores in community and host interactions. 3. Nat Rev Microbiol 18:152–163.

4. Hider RC, Kong X. 2010. Chemistry and biology of siderophores. Nat Prod Rep 27:637–657.

5. Schalk IJ. 2025. Bacterial siderophores: diversity, uptake pathways and applications. Nat Rev Microbiol 23:24–40.

6. Patel KD, Fisk MB, Gulick AM. 2024. Discovery, functional characterization, and structural studies of the NRPS-independent siderophore synthetases. Crit Rev Biochem Mol Biol 59:447–471.

7. Reitz ZL. 2024. Predicting metallophore structure and function through genome mining. Methods Enzymol 702:371–401.

8. Reitz ZL, Medema MH. 2022. Genome mining strategies for metallophore discovery. Curr Opin Biotechnol 77:102757.

9. Blin K, Shaw S, Vader L, Szenei J, Reitz ZL, Augustijn HE, Cediel-Becerra JDD, de Crécy-Lagard V, Koetsier RA, Williams SE, Cruz-Morales P, Wongwas S, Segurado Luchsinger AE, Biermann F, Korenskaia A, Zdouc MM, Meijer D, Terlouw BR, van der Hooft JJJ, Ziemert N, Helfrich EJN, Masschelein J, Corre C, Chevrette MG, van Wezel GP, Medema MH, Weber T. 2025. antiSMASH 8.0: extended gene cluster detection capabilities and analyses of chemistry, enzymology, and regulation. Nucleic Acids Res 53:W32–W38.

10. Skinnider MA, Johnston CW, Gunabalasingam M, Merwin NJ, Kieliszek AM, MacLellan RJ, Li H, Ranieri MRM, Webster ALH, Cao MPT, Pfeifle A, Spencer N, To QH, Wallace DP, Dejong CA, Magarvey NA. 2020. Comprehensive prediction of secondary metabolite structure and biological activity from microbial genome sequences. Nat Commun 11:6058.

11. Wu C, Medema MH, Läkamp RM, Zhang L, Dorrestein PC, Choi YH, Van Wezel GP. 2016. Leucanicidin and Endophenasides Result from Methyl-Rhamnosylation by the Same Tailoring Enzymes in *Kitasatospora* sp. MBT66. ACS Chem Biol 11:478–490.

12. Lazos O, Tosin M, Slusarczyk AL, Boakes S, Cortés J, Sidebottom PJ, Leadlay PF. 2010. Biosynthesis of the Putative Siderophore Erythrochelin Requires Unprecedented Crosstalk Between Separate Nonribosomal Peptide Gene Clusters. Chemistry & Biology 17:160–173.

13. Bosello M, Robbel L, Linne U, Xie X, Marahiel MA. 2011. Biosynthesis of the Siderophore Rhodochelin Requires the Coordinated Expression of Three Independent Gene Clusters in *Rhodococcus jostii* RHA1. J Am Chem Soc 133:4587–4595.

14. Baars O, Morel FMM, Perlman DH. 2014. ChelomEx: Isotope-Assisted Discovery of Metal Chelates in Complex Media Using High-Resolution LC-MS. Anal Chem 86:11298–11305.

15. Aron AT, Petras D, Schmid R, Gauglitz JM, Büttel I, Antelo L, Zhi H, Nuccio S-P, Saak CC, Malarney KP, Thines E, Dutton RJ, Aluwihare LI, Raffatellu M, Dorrestein PC. 2022. Native mass spectrometry-based metabolomics identifies metal-binding compounds. Nat Chem 14:100–109.

16. da Silva RR, Dorrestein PC, Quinn RA. 2015. Illuminating the dark matter in metabolomics. Proc Natl Acad Sci USA 112:12549–12550.

17. Gross H, Stockwell VO, Henkels MD, Nowak-Thompson B, Loper JE, Gerwick WH. 2007. The genomisotopic approach: a systematic method to isolate products of orphan biosynthetic gene clusters. Chem Biol 14:53–63.

18. McCaughey CS, van Santen JA, van der Hooft JJJ, Medema MH, Linington RG. 2022. An isotopic labeling approach linking natural products with biosynthetic gene clusters. Nat Chem Biol 18:295–304.

19. Wallace M, Cummings, Jr. DA, Roberts AG, Puri AW. 2023. A widespread methylotroph acyl-homoserine lactone synthase produces a new quorum sensing signal that regulates swarming in *Methylobacterium fujisawaense*. mBio 15:e01999–23.

20. Liebergesell TCE, Puri AW. 2024. Linking biosynthetic genes to natural products using inverse stable isotopic labeling (InverSIL). Methods Enzymol 702:215–227

21. Brady SF, Clardy J. 2005. Systematic Investigation of the *Escherichia coli* Metabolome for the Biosynthetic Origin of an Isocyanide Carbon Atom. Angew Chem Int Ed 44:7045–7048.

22. Brachmann AO, Forst S, Furgani GM, Fodor A, Bode HB. 2006. Xenofuranones A and B: Phenylpyruvate Dimers from *Xenorhabdus szentirmaii*. J Nat Prod 69:1830–1832.

23. Reitz ZL, Sandy M, Butler A. 2017. Biosynthetic considerations of triscatechol siderophores framed on serine and threonine macrolactone scaffolds. Metallomics 9:824–839.

24. Loomis LD, Raymond KN. 1991. Solution equilibria of enterobactin and metal-enterobactin complexes. Inorg Chem 30:906–911.

25. May JJ, Wendrich TM, Marahiel MA. 2001. The *dhb* Operon of *Bacillus subtilis* Encodes the Biosynthetic Template for the Catecholic Siderophore 2,3-Dihydroxybenzoate-Glycine-Threonine Trimeric Ester Bacillibactin. J Biol Chem 276:7209–7217.

26. Pfleger BF, Kim Y, Nusca TD, Maltseva N, Lee JY, Rath CM, Scaglione JB, Janes BK, Anderson EC, Bergman NH, Hanna PC, Joachimiak A, Sherman DH. 2008. Structural and functional analysis of AsbF: Origin of the stealth 3,4-dihydroxybenzoic acid subunit for petrobactin biosynthesis. Proc Natl Acad Sci USA 105:17133–17138.

27. Fox DT, Hotta K, Kim C-Y, Koppisch AT. 2008. The Missing Link in Petrobactin Biosynthesis: *asbF* Encodes a (−)-3-Dehydroshikimate Dehydratase. Biochemistry 47:12251–12253.

28. Liebergesell TCE, Murdock EG, Puri AW. 2024. Detection of Inverse Stable Isotopic Labeling in Untargeted Metabolomic Data. Anal Chem 96:16330–16337.

29. Robes JMD, Liebergesell TCE, Beals DG, Yu X, Brazelton WJ, Puri AW. 2025. Inverse stable isotope probing–metabolomics (InverSIP) identifies an iron acquisition system in a methane-oxidizing bacterial community. Proc Natl Acad Sci USA 122:e2507323122.

30. Chistoserdova L. 2015. Methylotrophs in natural habitats: current insights through metagenomics. Appl Microbiol Biotechnol 99:5763–5779.

31. Zheng Y, Huang J, Zhao F, Chistoserdova L. 2018. Physiological Effect of XoxG on Lanthanide-Dependent Methanotrophy. mBio 9:e02430–17.

32. Esmaeel Q, Pupin M, Kieu NP, Chataigné G, Béchet M, Deravel J, Krier F, Höfte M, Jacques P, Leclère V. 2016. Burkholderia genome mining for nonribosomal peptide synthetases reveals a great potential for novel siderophores and lipopeptides synthesis. Microbiologyopen 5:512–526.

33. Baars O, Zhang X, Morel FMM, Seyedsayamdost MR. 2015. The Siderophore Metabolome of *Azotobacter vinelandii*. Appl Environ Microbiol 82:27–39.

34. Beck DAC, McTaggart TL, Setboonsarng U, Vorobev A, Kalyuzhnaya MG, Ivanova N, Goodwin L, Woyke T, Lidstrom ME, Chistoserdova L. 2014. The expanded diversity of Methylophilaceae from Lake Washington through cultivation and genomic sequencing of novel ecotypes. PLoS ONE 9:e102458.

35. Fedorov DN, Doronina NV, Trotsenko YA. 2011. Phytosymbiosis of aerobic methylobacteria: New facts and views. Microbiology 80:443–454.

36. Kovaleva J, Degener JE, van der Mei HC. 2014. *Methylobacterium* and Its Role in Health Care-Associated Infection. J Clin Microbiol 52:1317–1321.

37. Zytnick AM, Gutenthaler-Tietze SM, Aron AT, Reitz ZL, Phi MT, Good NM, Petras D, Daumann LJ, Martinez-Gomez NC. 2024. Identification and characterization of a small-molecule metallophore involved in lanthanide metabolism. Proc Natl Acad Sci USA 121:e2322096121.

38. Baars O, Morel FMM, Zhang X. 2018. The purple non-sulfur bacterium *Rhodopseudomonas palustris* produces novel petrobactin-related siderophores under aerobic and anaerobic conditions. Environ Microbiol 20:1667–1676.

39. Batista BB, Santos RERDS, Ricci-Azevedo R, Da Silva Neto JF. 2019. Production and Uptake of Distinct Endogenous Catecholate-Type Siderophores Are Required for Iron Acquisition and Virulence in *Chromobacterium violaceum*. Infect Immun 87:e00577–19.

40. Butler A, Jelowicki AM, Ogasawara HA, Reitz ZL, Stow PR, Thomsen E. 2023. Mining elements of siderophore chirality encoded in microbial genomes. FEBS Lett 597:134–140.

41. Sandy M, Butler A. 2011. Chrysobactin Siderophores Produced by *Dickeya chrysanthemi* EC16. J Nat Prod 74:1207–1212.

42. Navarro-Torre S, Carro L, Rodríguez-Llorente ID, Pajuelo E, Caviedes MÁ, Igual JM, Redondo-Gómez S, Camacho M, Klenk H-P, Montero-Calasanz MDC. 2018. *Kushneria phyllosphaerae* sp. nov. and *Kushneria endophytica* sp. nov., plant growth promoting endophytes isolated from the halophyte plant *Arthrocnemum macrostachyum*. Int J Syst Evol Microbiol 68:2800–2806.

43. Meinzer M, Ahmad N, Nielsen BL. 2023. Halophilic Plant-Associated Bacteria with Plant-Growth-Promoting Potential. Microorganisms 11:2910.

44. Peterson T, Nielands JB. 1979. Revised structure of a catecholamide spermidine siderophore. Tetrahedron Lett 20:4805–4808.

45. Tait GH. 1975. The identification and biosynthesis of siderochromes formed by *Micrococcus denitrificans*. Biochem J 146:191–204.

46. Wang H, Fewer DP, Holm L, Rouhiainen L, Sivonen K. 2014. Atlas of nonribosomal peptide and polyketide biosynthetic pathways reveals common occurrence of nonmodular enzymes. Proc Natl Acad Sci USA 111:9259–9264.

47. González Carreró MI, Sangari FJ, Agüero J, García Lobo JM. 2002. *Brucella abortus* strain 2308 produces brucebactin, a highly efficient catecholic siderophore. Microbiology 148:353–360.

48. Vogel CM, Potthoff DB, Schäfer M, Barandun N, Vorholt JA. 2021. Protective role of the *Arabidopsis* leaf microbiota against a bacterial pathogen. Nat Microbiol 6:1537–1548.

49. Dimise EJ, Widboom PF, Bruner SD. 2008. Structure elucidation and biosynthesis of fuscachelins, peptide siderophores from the moderate thermophile *Thermobifida fusca*. Proc Natl Acad Sci USA 105:15311–15316.

50. Boiteau RM, Fansler SJ, Farris Y, Shaw JB, Koppenaal DW, Pasa-Tolic L, Jansson JK. 2019. Siderophore profiling of co-habitating soil bacteria by ultra-high resolution mass spectrometry. Metallomics 11:166–175.

51. Gubbens J, Wu C, Zhu H, Filippov DV, Florea BI, Rigali S, Overkleeft HS, Van Wezel GP. 2017. Intertwined Precursor Supply during Biosynthesis of the Catecholate–Hydroxamate Siderophores Qinichelins in Streptomyces sp. MBT76. ACS Chem Biol 12:2756–2766.

52. McRose DL, Seyedsayamdost MR, Morel FMM. 2018. Multiple siderophores: bug or feature? J Biol Inorg Chem 23:983–993.

53. Ventosa A, Quesada E, Rodriguez-Valera F, Ruiz-Berraquero F, Ramos-Cormenzana A. 1982. Numerical Taxonomy of Moderately Halophilic Gram-negative Rods. Microbiology 128:1959–1968.

54. Hirakawa H, Schaefer AL, Greenberg EP, Harwood CS. 2012. Anaerobic *p*-Coumarate Degradation by *Rhodopseudomonas palustris* and Identification of CouR, a MarR Repressor Protein That Binds *p*-Coumaroyl Coenzyme A. J Bacteriol 194:1960–1967.

55. Markowitz VM, Chen I-MA, Palaniappan K, Chu K, Szeto E, Grechkin Y, Ratner A, Jacob B, Huang J, Williams P, Huntemann M, Anderson I, Mavromatis K, Ivanova NN, Kyrpides NC. 2012. IMG: the Integrated Microbial Genomes database and comparative analysis system. Nucleic Acids Res 40:D115–122.

56. van den Belt M, Gilchrist C, Booth TJ, Chooi Y-H, Medema MH, Alanjary M. 2023. CAGECAT: The CompArative GEne Cluster Analysis Toolbox for rapid search and visualisation of homologous gene clusters. BMC Bioinform 24:181.

