## Supplemental Info for "Inverse stable isotope labeling (InverSIL) links predicted catecholate siderophore gene clusters to their products in diverse bacteria"

**TABLE OF CONTENTS**

SUPPLEMENTARY FIGURES ... 3
Figure S1. ... 3
Figure S2 ... 4
Figure S3 ... 5
Figure S4 ... 6
Figure S5 ... 7
Figure S6 ... 8
Figure S7 ... 9
Figure S8 ... 10
Figure S9 ... 11
Figure S10 ... 12
Figure S11 ... 13
Figure S12 ... 14
Figure S13 ... 15
Figure S14 ... 16
Figure S15 ... 17
Figure S16 ... 18
Figure S17 ... 19
SUPPLEMENTARY TABLES ... 20
Table S1 ... 20
Table S2 ... 21
Table S3 ... 22
Table S4 ... 23
Table S5 ... 24
Table S6 ... 25
Table S7 ... 26
Table S8 ... 28
Table S9 ... 29
Table S10 ... 30
Table S11 ... 31
REFERENCES ... 32

### SUPPLEMENTARY FIGURES

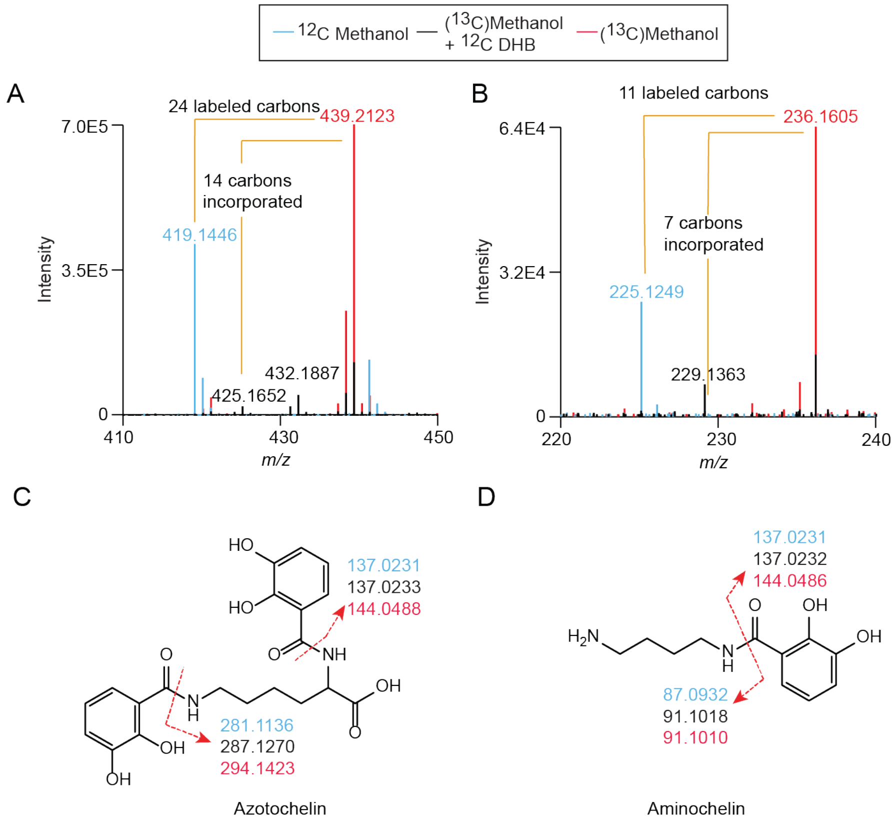


**Figure S1.** *Methylophilus* sp. strain 5 InverSIL showing incorporation of 2,3-DHB into (A) azotochelin and (B) aminochelin. (C) Structure of azotochelin with fragmentation data from different growth conditions. (D) Structure of aminochelin with fragmentation data from different growth conditions.

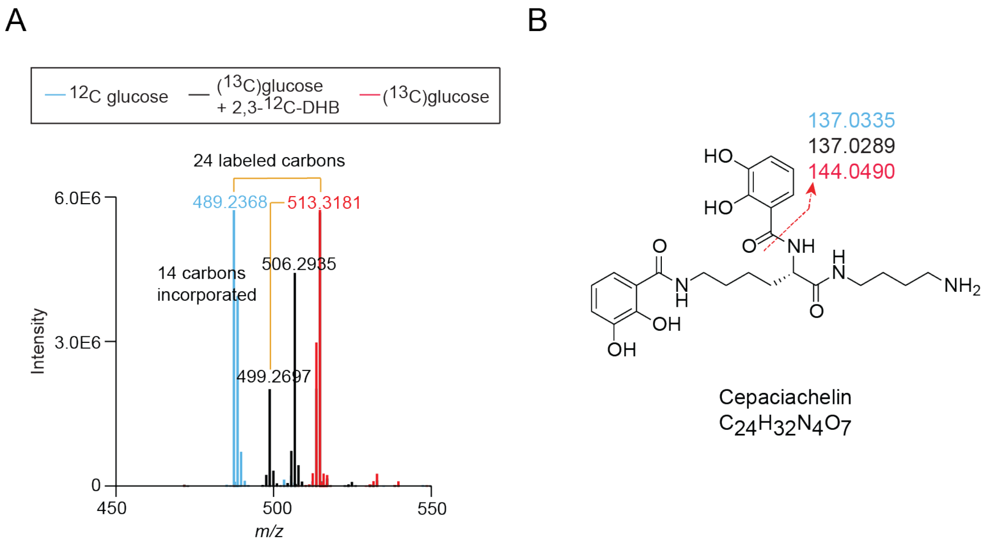


**Figure S2**. Using InverSIL to determine the structure of the chromobactin BGC product from *C. violaceum* CV017. (A) Overlayed mass spectra of *C. violaceum* CV017 supernatant extract showing incorporation of two 2,3-DHB units into a metabolite with the same high-resolution mass and carbon count as cepaciachelin. (B) Structure of chromobactin (cepaciachelin) showing MS2 fragments from different inverSIL conditions. The colors match the growth conditions indicated in panel A.

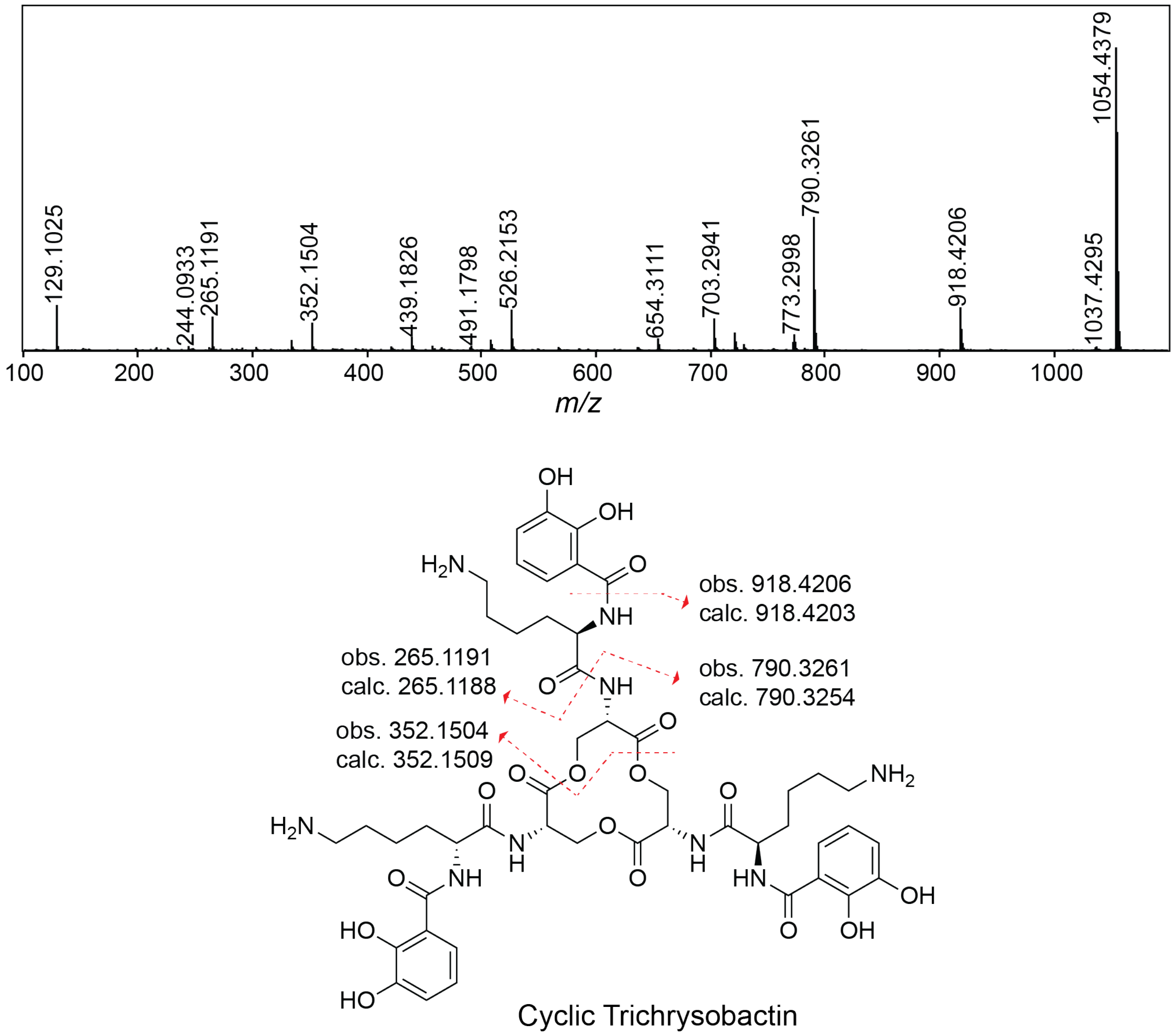


Cyclic Trichrysobactin

**Figure S3**. Fragmentation of cyclic trichrysobactin in *C. violaceum* CV017. Key fragments are highlighted and compared to Sandy and Butler (2012) (1).

A

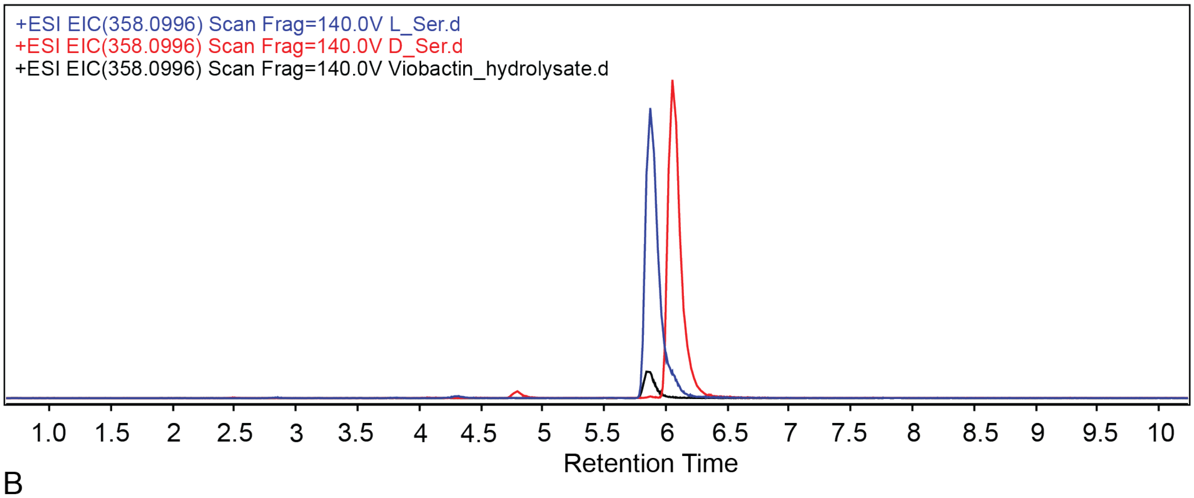


B

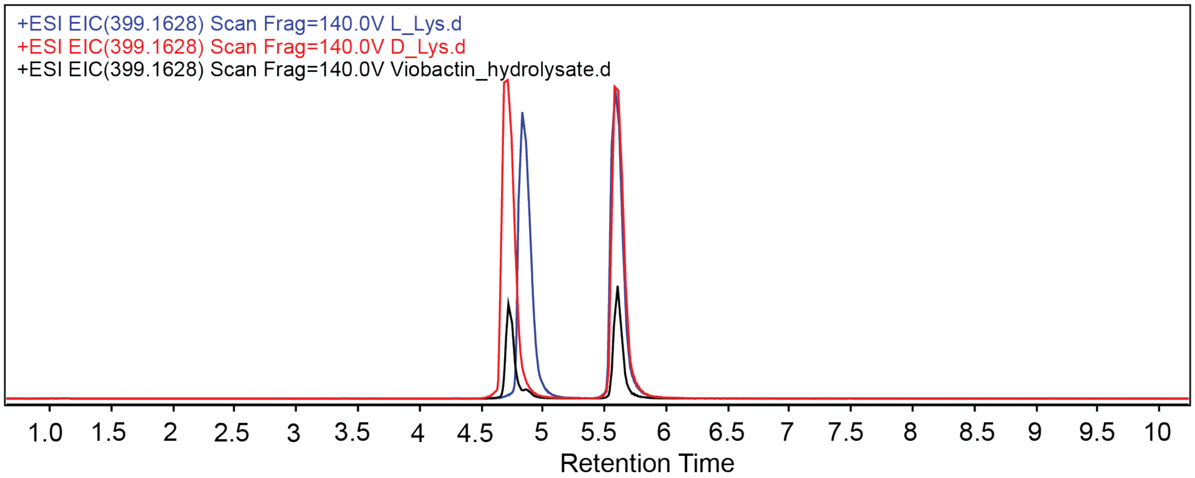


**Figure S4**. Amino acid stereochemistry determination of viobactin using Marfey's analysis. (A) Extracted ion chromatogram for m/z 358.0996 corresponding to $[M+H]^+$ of Marfey's derivatized serine. Blue (Marfey's derivatized L-serine), red (Marfey's derivatized D-serine), and black (Marfey's derivatized cellulochelin B hydrolysate). Mass tolerance < 5ppm. (B) Extracted ion chromatogram for m/z 399.1628 corresponding to $[M+H]^+$ of Marfey's derivatized ornithine. Blue (Marfey's derivatized L-ornithine), red (Marfey's derivatized D-ornithine), and black (Marfey's derivatized cellulochelin B hydrolysate)Mass tolerance < 5ppm. Related to Table S4.

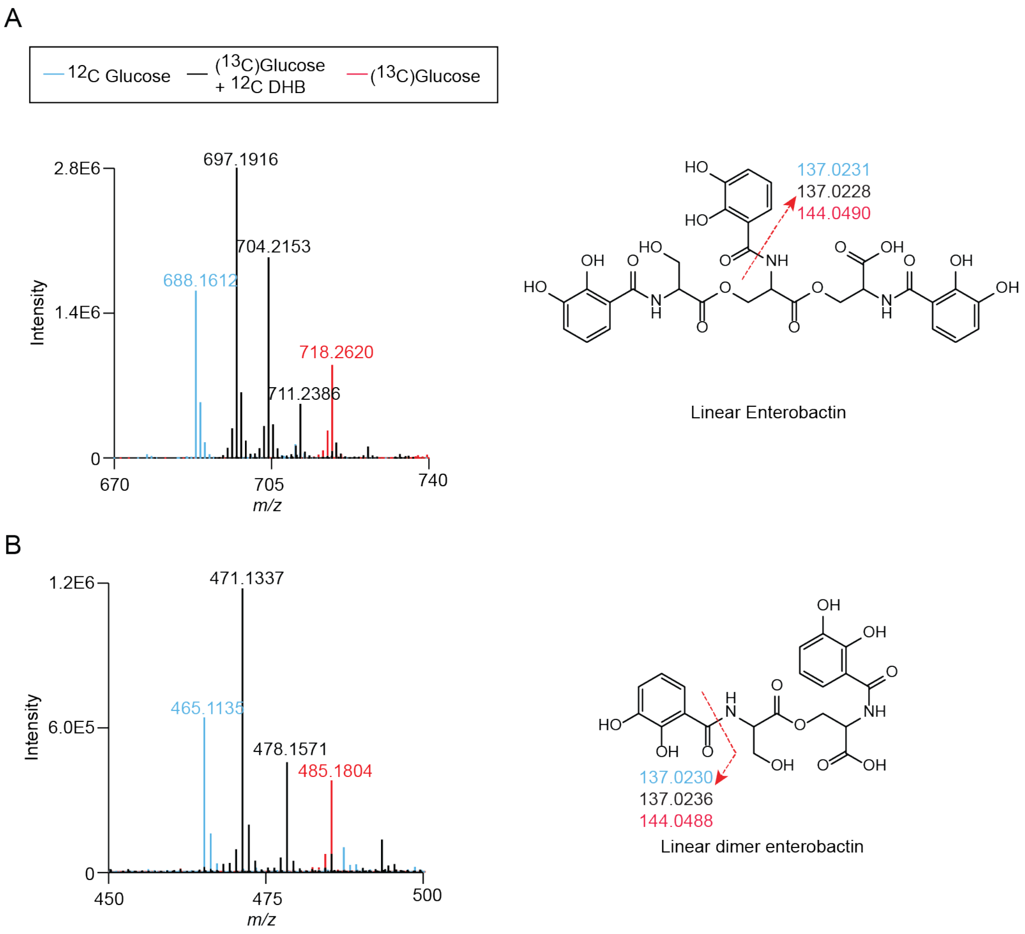


**Figure S5**. (A, B) Other enterobactin forms in *Kushneria konosiri* JCM16805 detected using InverSIL.

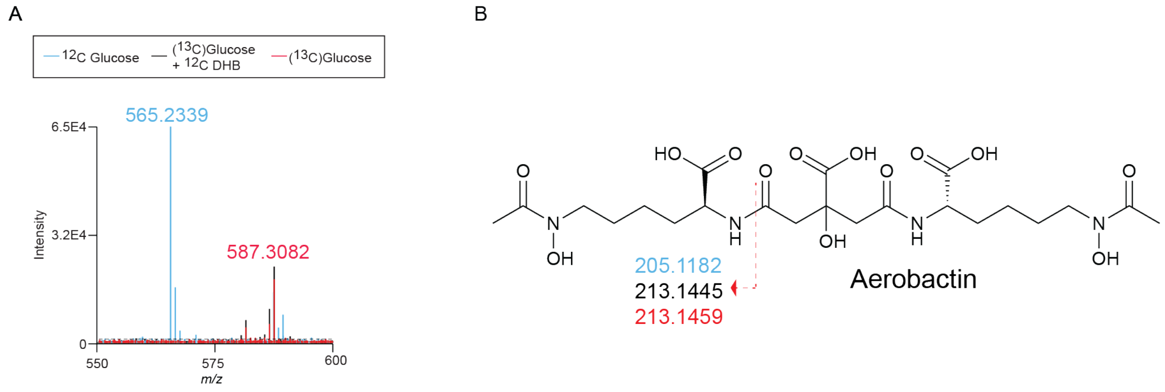


**Figure S6**. Mass spectrum of aerobactin in *K. konosiri* JCM 16805 grown in $^{12}$C glucose (cyan), ($^{13}$C)glucose (red), ($^{13}$C)glucose + 2,3DHB (black). (B) Structure and MS2 fragmentation analysis of aerobactin.

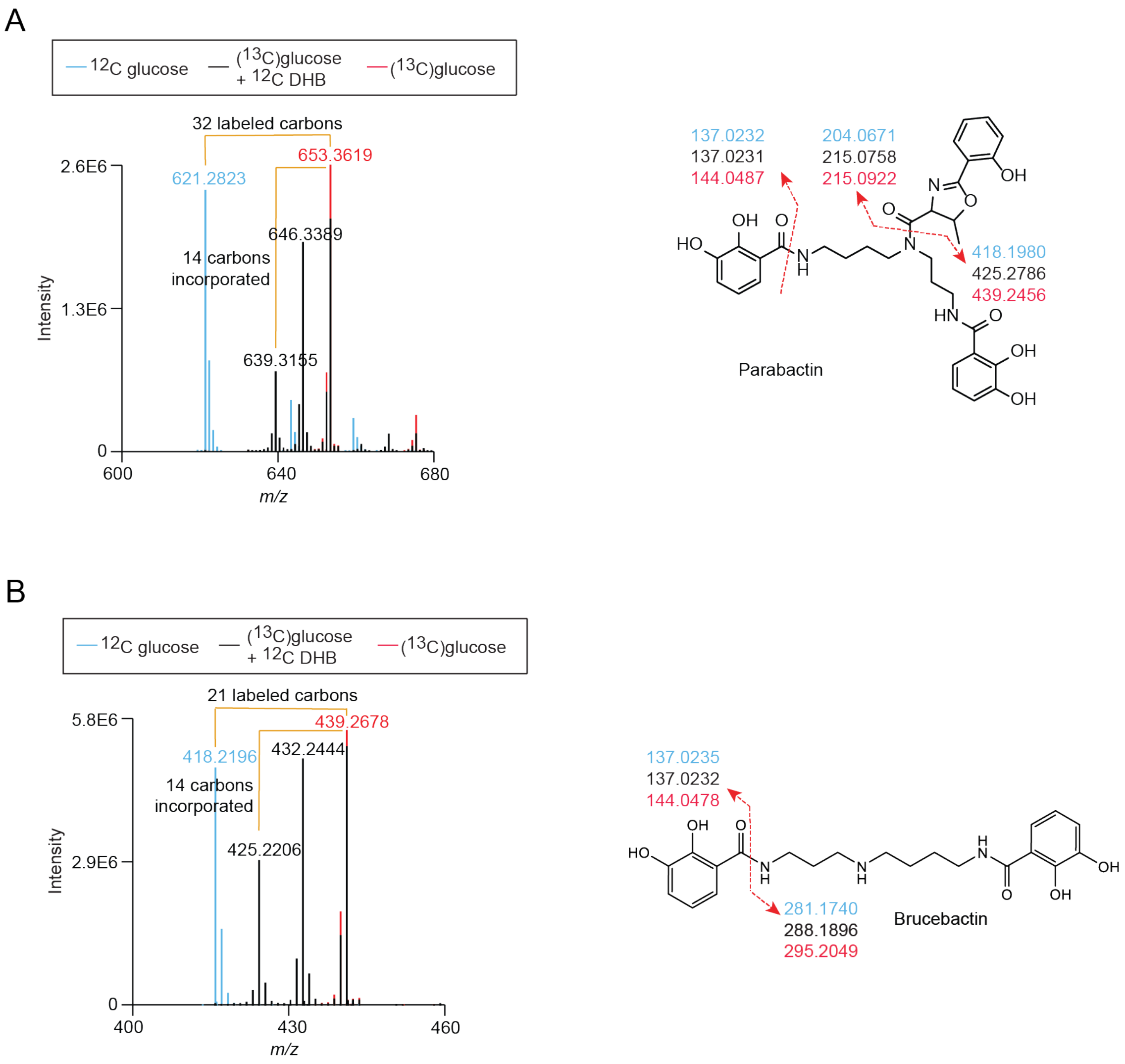


**Figure S7**. Detection and inverse labeling of (A) parabactin and (B) brucebactin in *Paracoccus denitrificans* PD1222.

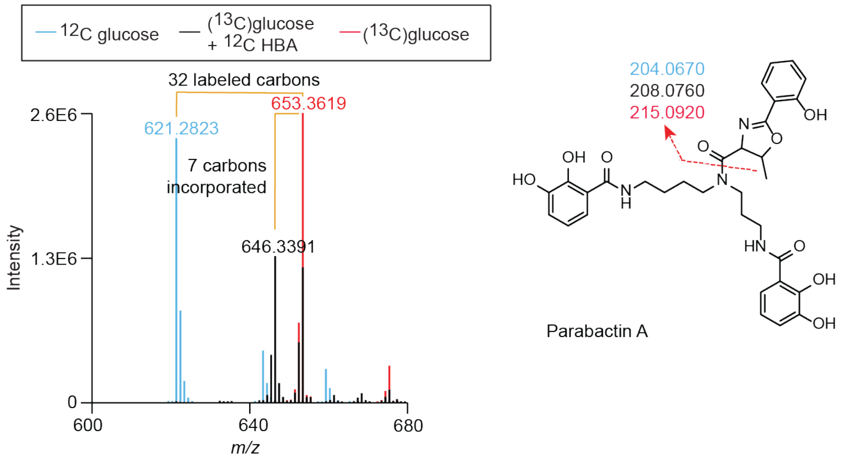


**Figure S8**. Inverse labeling of 2-hydroxybenzoic acid in parabactin.

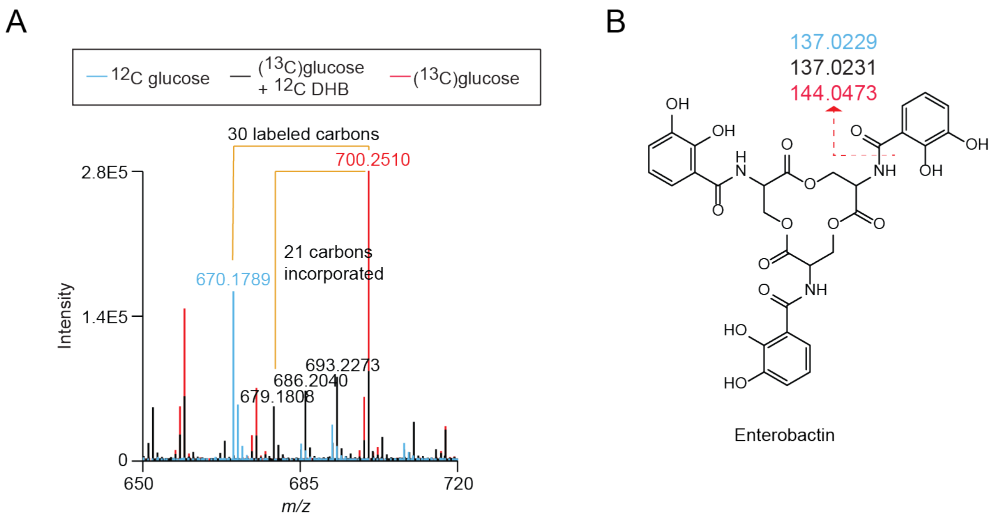


**Figure S9**. Inverse labeling of 2,3-DHB in enterobactin from *Paracoccus denitrificans* PD1222.

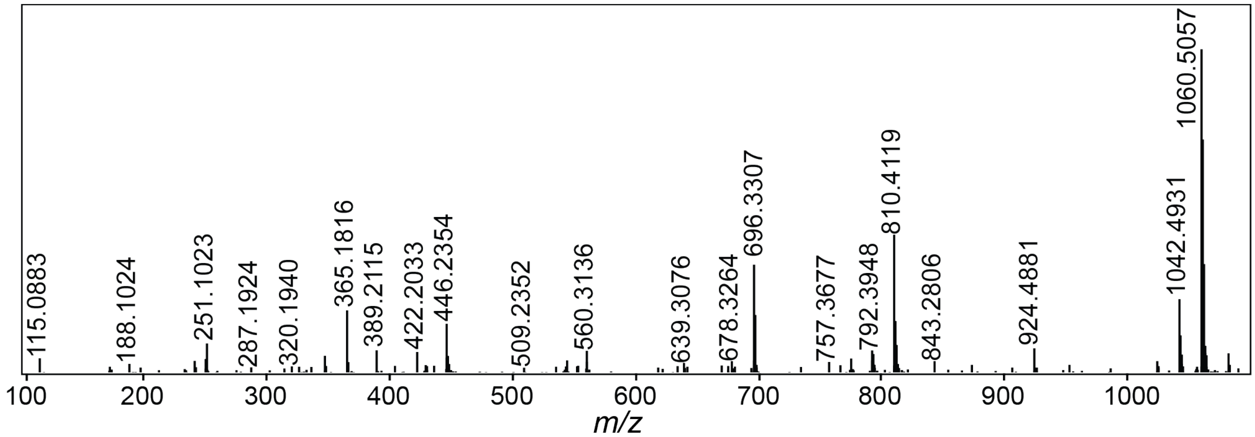


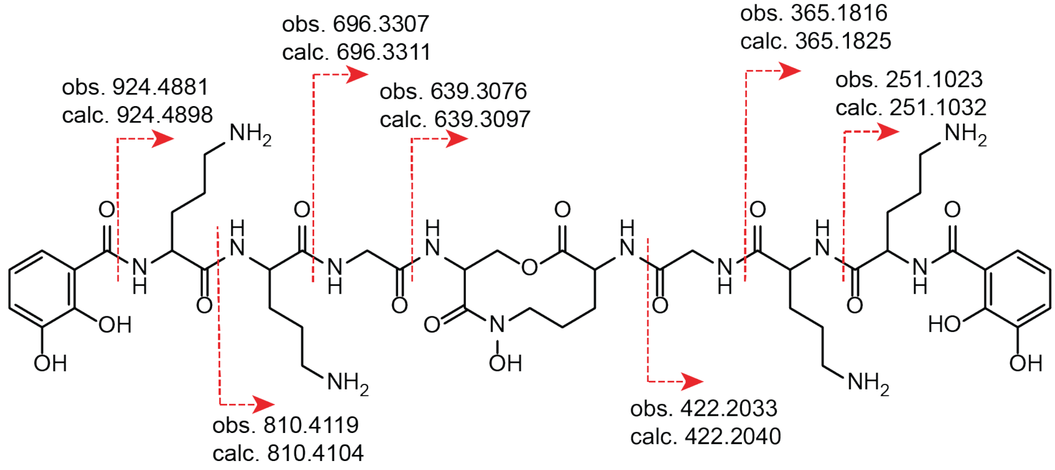


Cellulochelin A

**Figure S10**. MS2 fragmentation of cellulochelin A.

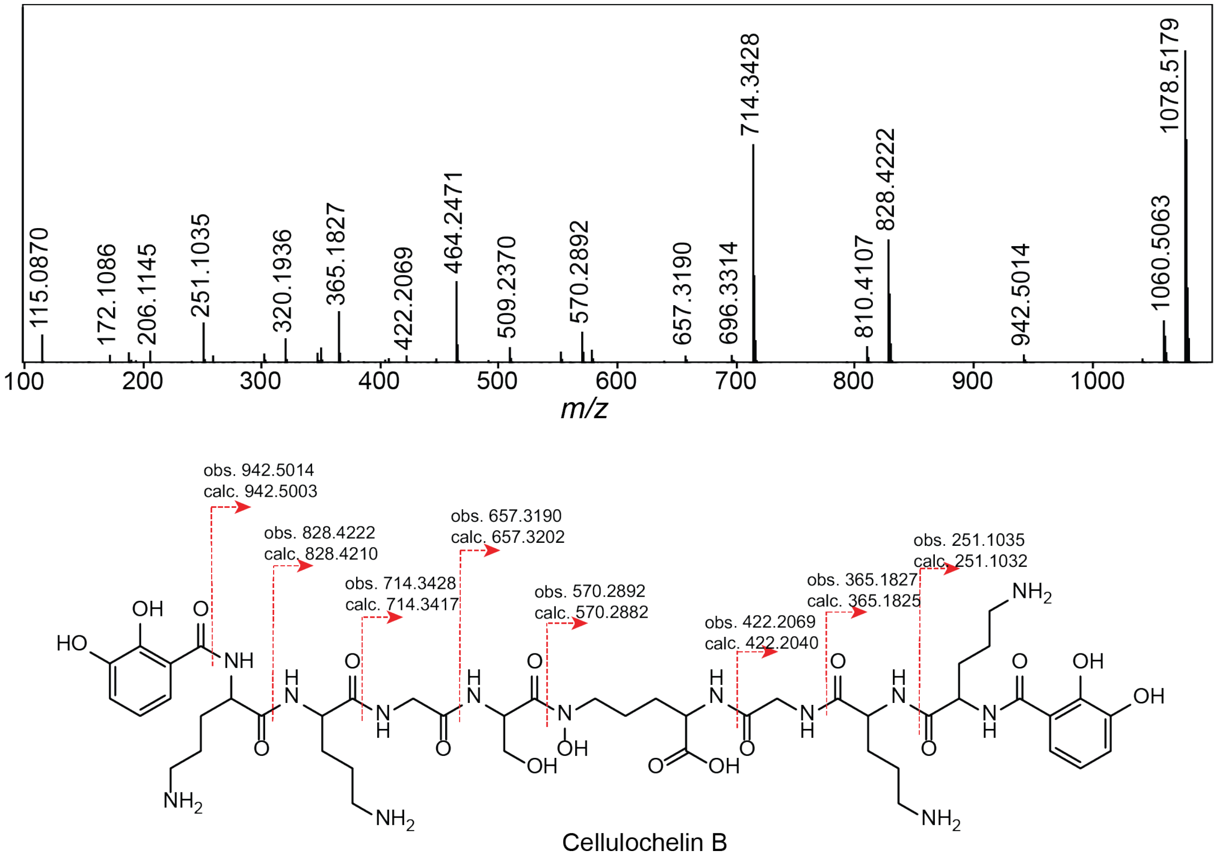


**Figure S11**. MS2 fragmentation of cellulochelin B.

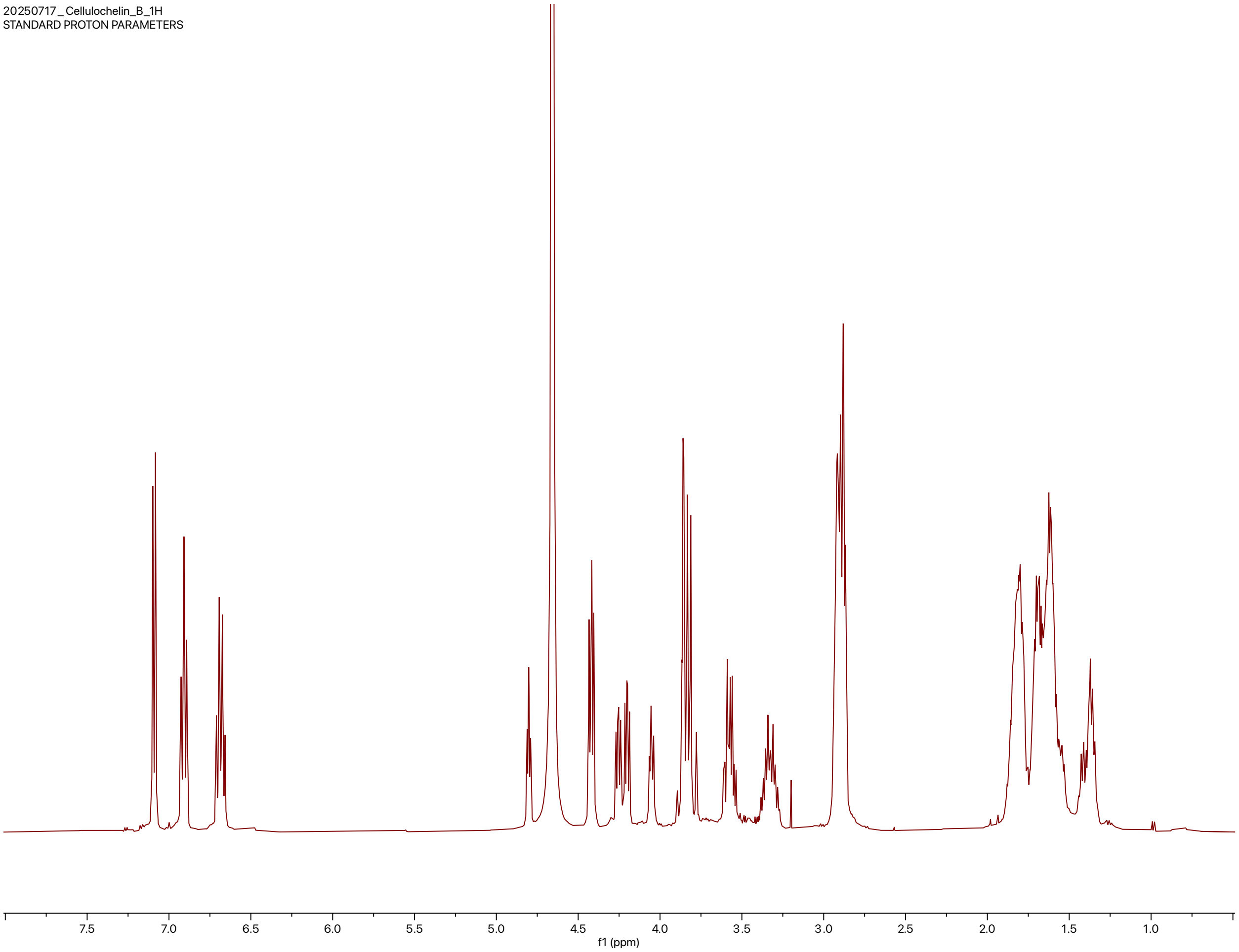


**Figure S12**. [1]H NMR spectrum of cellulochelin B in $D_2O$ (500 MHz).

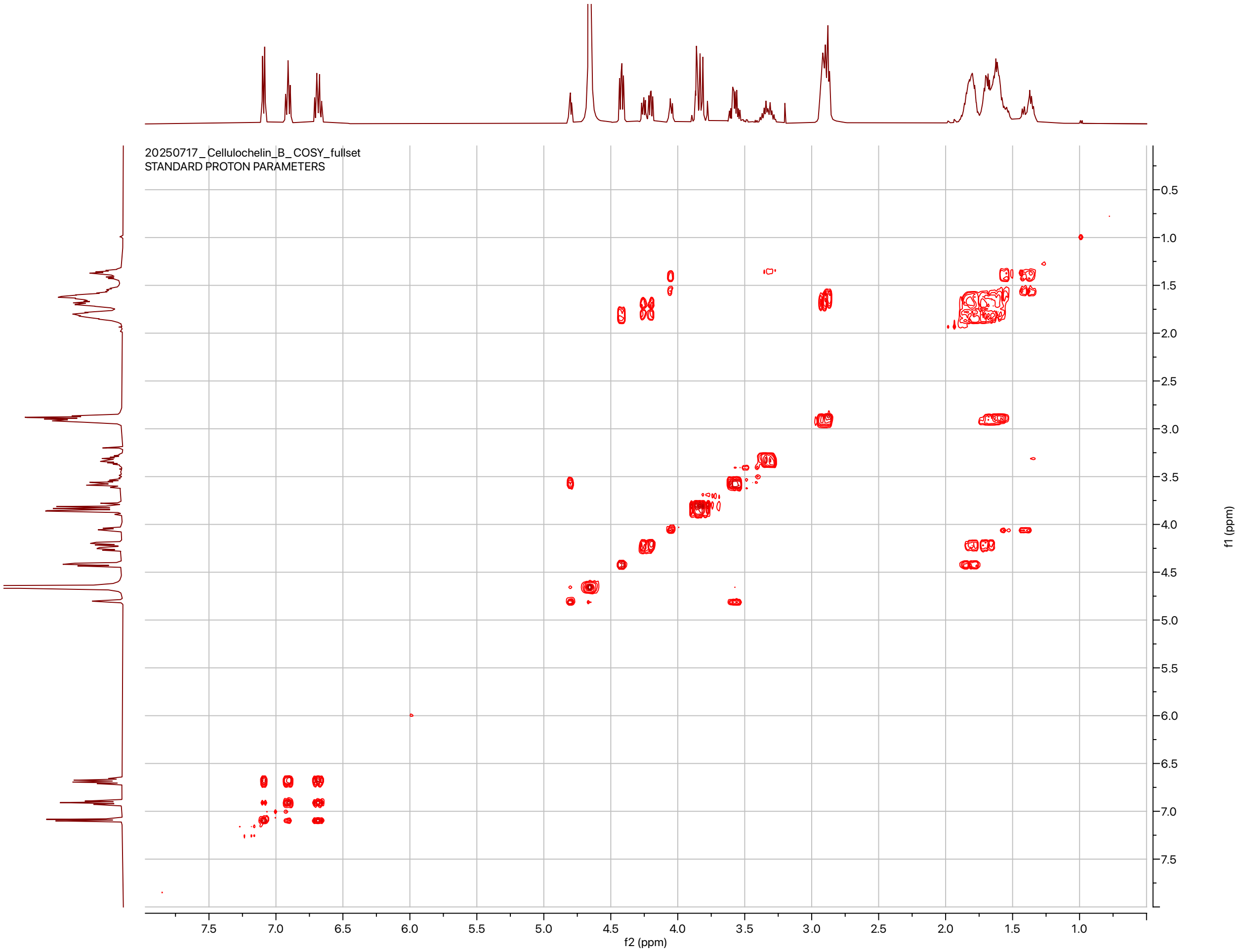


**Figure S13**. COSY NMR spectrum of cellulochelin B in $D_2O$ (500 MHz)**.**

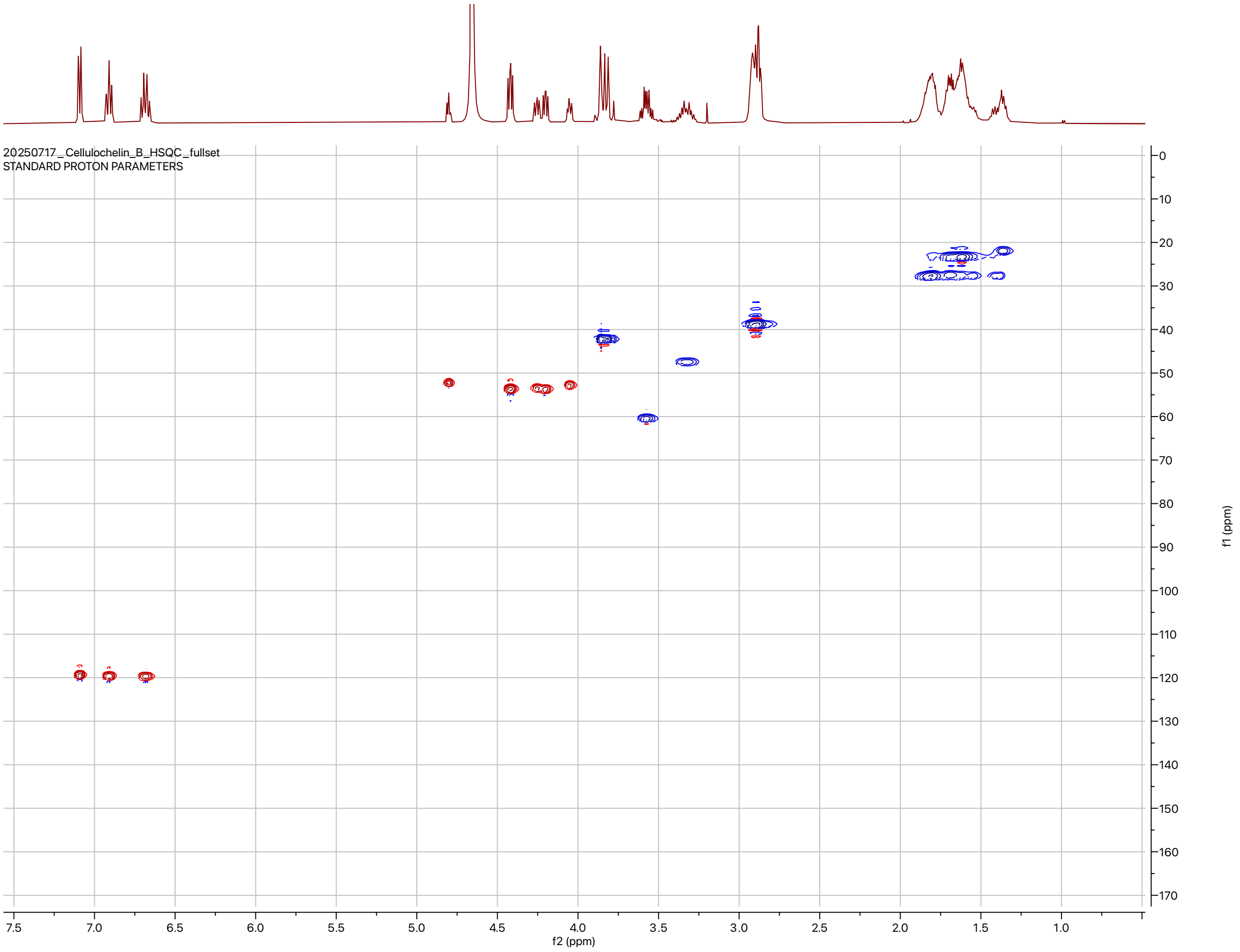


**Figure S14**. gHSQC spectrum of cellulochelin B in $D_2O$ (500 MHz)**.**

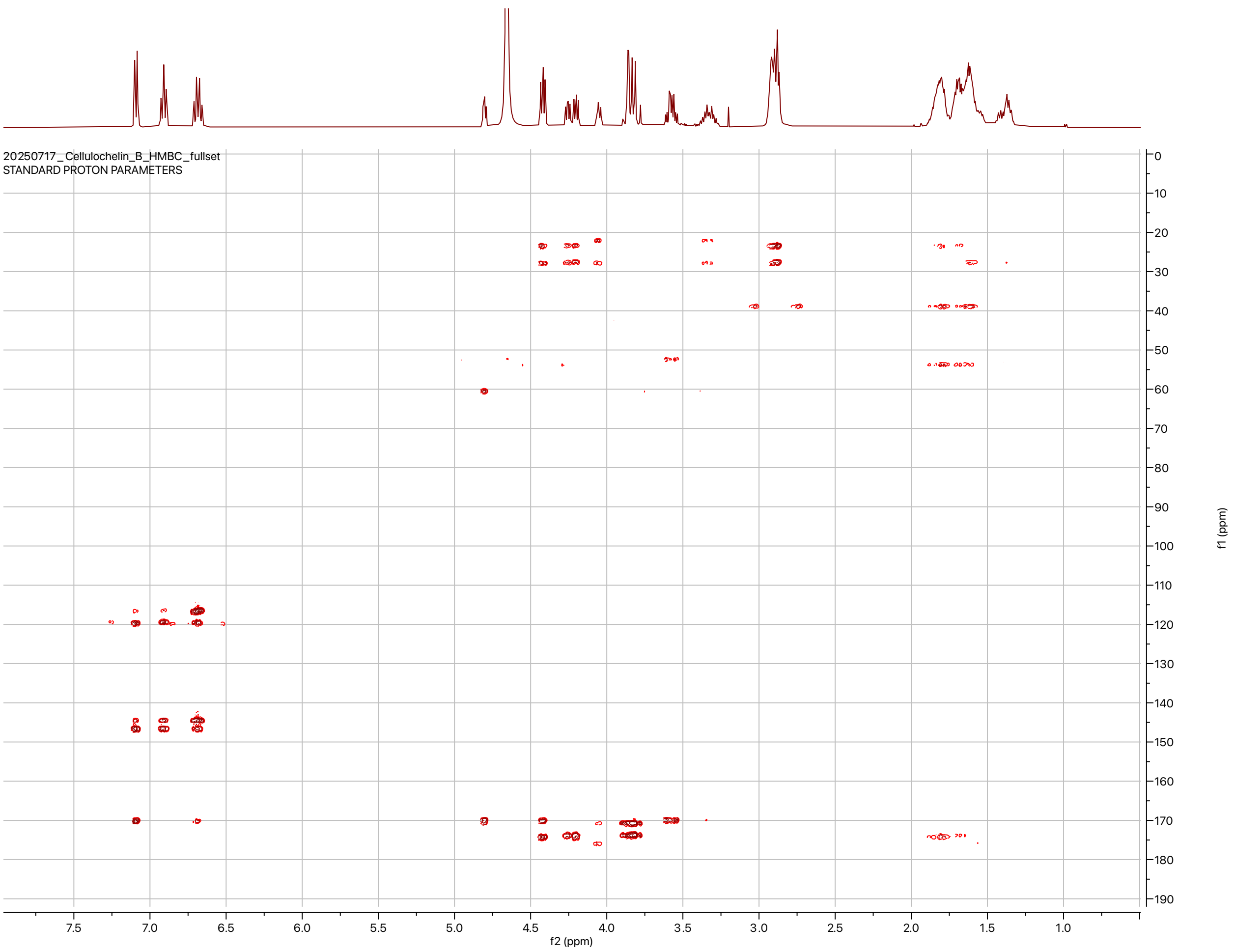


**Figure S15**. HMBC NMR spectrum of cellulochelin B in $D_2O$ (500 MHz).

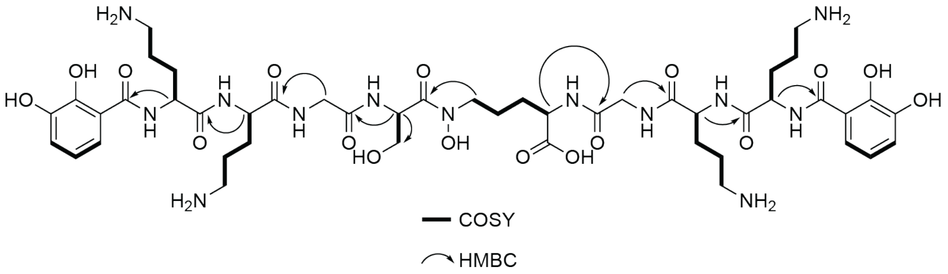


**Figure S16**. COSY and HMBC NMR correlations in cellulochelin B in $D_2O$ (500 MHz)**.**

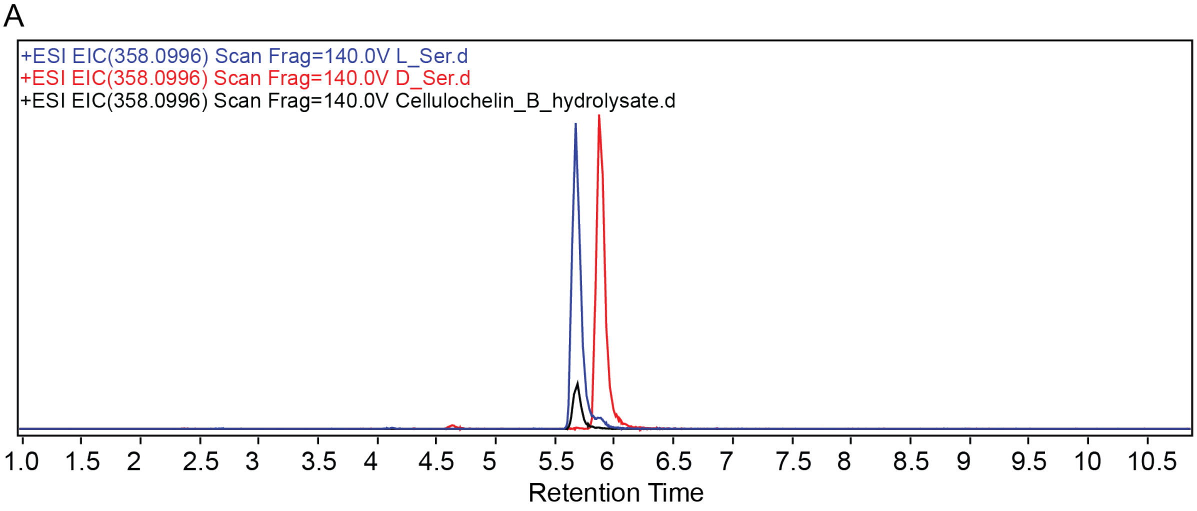


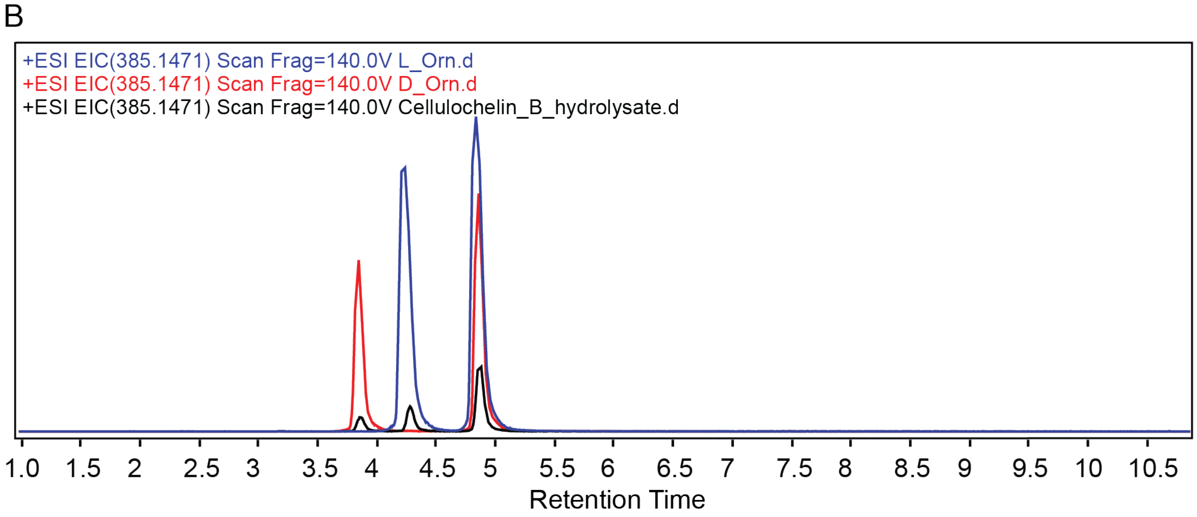


**Figure S17**. Amino acid stereochemistry determination of cellulochelin B using Marfey's analysis. (A) Extracted ion chromatogram for m/z 358.0996 corresponding to $[M+H]^+$ of Marfey's derivatized serine. Blue (Marfey's derivatized L-serine), red (Marfey's derivatized D-serine), and black (Marfey's derivatized cellulochelin B hydrolysate). Mass tolerance < 5ppm. (B) Extracted ion chromatogram for m/z 385.1471 corresponding to $[M+H]^+$ of Marfey's derivatized ornithine. Blue (Marfey's derivatized L-ornithine), red (Marfey's derivatized D-ornithine), and black (Marfey's derivatized cellulochelin B hydrolysate) Mass tolerance < 5ppm. Related to Table S8.

### SUPPLEMENTARY TABLES

**Table S1.** Cepaciachelin MS/MS fragmentation comparison between crude extracts from *B. ambifaria* BAA244, *Methylophilus* sp. strain 5, and *C. violaceum* CV017.

| *Burkholderia ambifaria* BAA244 | | *Methylophilus* sp. strain 5 | | *Chromobacterium violaceum* CV017 | |
|---|---|---|---|---|---|
| *m/z* | Intensity | *m/z* | Intensity | *m/z* | Intensity |
| 84.0903 | 3.5E+06 | 84.0805 | 2.5E+06 | 84.0809 | 3.5E+05 |
| 72.0896 | 2.6E+06 | 72.0804 | 1.8E+06 | 72.0809 | 2.4E+05 |
| 89.1171 | 1.7E+06 | 89.1069 | 1.2E+06 | 89.1074 | 1.4E+05 |
| 220.1124 | 1.5E+06 | 220.0967 | 1.1E+06 | 220.097 | 1.3E+05 |
| 353.2379 | 7.3E+05 | 353.2185 | 5.7E+05 | 353.2186 | 6.1E+04 |
| 217.2178 | 2.8E+05 | 217.2023 | 1.7E+05 | 217.2025 | 2.4E+04 |
| 85.0937 | 2.4E+05 | 85.0837 | 1.5E+05 | 221.1004 | 2.2E+04 |
| 221.1158 | 2.4E+05 | 129.1021 | 1.1E+05 | 85.0842 | 2.1E+04 |
| 129.1141 | 1.7E+05 | 354.2218 | 1.1E+05 | 354.2218 | 2.0E+04 |
| 137.0355 | 1.7E+05 | 137.0231 | 9.9E+04 | 137.0234 | 1.6E+04 |

**Table S2.** Rhodopetrobactin B MS/MS fragmentation comparison between crude extracts from *R. palustris* CGA009 and *M. extorquens* PA1.

| *Rhodopseudomonas palustris* CGA009 | | *Methylorubrum extorquens* PA1 | |
|---|---|---|---|
| *m/z* | Intensity | *m/z* | Intensity |
| 338.2078 | 1.0E+05 | 338.2084 | 1.2E+04 |
| 202.1913 | 1.0E+05 | 202.192 | 1.2E+04 |
| 677.3871 | 6.4E+04 | 677.3878 | 8.1E+03 |
| 541.3709 | 5.7E+04 | 541.3713 | 7.5E+03 |
| 312.1919 | 4.0E+04 | 312.1924 | 5.0E+03 |
| 185.1649 | 3.7E+04 | 185.1651 | 4.2E+03 |
| 321.1813 | 3.4E+04 | 321.182 | 3.8E+03 |
| 678.3909 | 2.7E+04 | 678.3914 | 2.7E+03 |
| 208.0971 | 2.2E+04 | 358.1978 | 2.6E+03 |
| 482.3345 | 2.1E+04 | 208.0979 | 2.4E+03 |

**Table S3**. Top 10 MS2 fragments of viobactin (cyclic trichrysobactin) from *C. violaceum* CV017

| *m/z* | Intensity |
|---|---|
| 1054.438 | 1.5E+05 |
| 790.3261 | 6.5E+04 |
| 1056.443 | 3.9E+04 |
| 791.3293 | 3.0E+04 |
| 129.1025 | 2.2E+04 |
| 918.4206 | 2.1E+04 |
| 526.2153 | 2.0E+04 |
| 265.1191 | 1.6E+04 |
| 703.2941 | 1.5E+04 |
| 352.1504 | 1.3E+04 |

**Table S4**. Advanced Marfey's analysis of viobactin. Related to Figure S4.

| Amino Acid | Analyte $t_r$ (min) | L-Standard $t_r$ (min) | D-Standard $t_r$ (min) |
|---|---|---|---|
| Serine | 5.84 | 5.87 | 6.05 |
| Lysine | 4.72 | 4.83 | 4.72 |

**Table S5**. Enterobactin MS/MS fragmentation comparison between crude extracts from *E. coli* MG1655, *K. konosiri* JCM16805, and *P. denitrificans* PD1222.

| *Escherichia coli* MG1655 | | *Kushneria konosiri* JCM16805 | | *Paracoccus denitrificans* PD1222 | |
|---|---|---|---|---|---|
| *m/z* | Intensity | *m/z* | Intensity | *m/z* | Intensity |
| 224.0552 | 2.3E+05 | 224.0552 | 2.0E+05 | 224.0555 | 3.8E+05 |
| 225.0584 | 2.9E+04 | 178.0493 | 2.9E+04 | 225.0585 | 4.9E+04 |
| 206.0446 | 2.7E+04 | 137.0228 | 2.8E+04 | 447.103 | 1.1E+04 |
| 137.0233 | 1.7E+04 | 225.0581 | 2.5E+04 | 206.0444 | 9.9E+03 |
| 447.1027 | 1.0E+04 | 150.0546 | 8.9E+03 | 311.0869 | 9.6E+03 |
| 196.06 | 7.9E+03 | 206.0445 | 7.3E+03 | 178.0498 | 8.7E+03 |
| 178.0495 | 6.5E+03 | 157.0608 | 4.2E+03 | 196.0606 | 7.6E+03 |
| 157.0606 | 5.4E+03 | 179.0528 | 4.1E+03 | 137.0234 | 7.3E+03 |
| 226.0591 | 4.4E+03 | 162.0553 | 3.7E+03 | 226.0596 | 6.8E+03 |
| 293.0765 | 4.3E+03 | 60.0439 | 3.1E+03 | 448.1067 | 4.1E+03 |

**Table S6**. Top 10 MS2 fragments of aerobactin from *K. konosiri* JCM16805

| *m/z* | Intensity |
|---|---|
| 205.1183 | 9.6E+03 |
| 100.0752 | 8.4E+03 |
| 142.0866 | 1.9E+03 |
| 159.1127 | 1.6E+03 |
| 273.1066 | 1.5E+03 |
| 145.0983 | 1.0E+03 |
| 301.1015 | 9.9E+02 |
| 315.1184 | 9.2E+02 |
| 206.1213 | 9.0E+02 |
| 128.0708 | 8.8E+02 |

**Table S7**. Summary of $^{1}H$ NMR data (δ in ppm) and $^{13}C$ NMR data (δ in ppm) for cellulochelin B in $D_2O$.

| Residue | Position | δC | δH, multiplicity (J in Hz) |
|---|---|---|---|
| 2,3-Dihydroxy benzoic acid (1) | C=O | 170 | - |
| | qC | 115.1 | - |
| | C | 146.4 | - |
| | C | 144.3 | - |
| | CH | 119.6 | 6.91, t (8.65) |
| | CH | 119.6 | 6.68, d (8.07) |
| | CH | 119.3 | 7.08, m |
| D-Ornithine (1) | α-CH | 53.6 | 4.42, dd (5.80, 8.74) |
| | β-CH2 | 27.7 | 1.85, m |
| | γ-CH2 | 23.8 | 1.65, m |
| | δ-CH2 | 38.6 | 2.88, m |
| | C=O | 174.1 | - |
| L-Ornithine (1) | α-CH | 53.4 | 4.25, dd (5.63, 8.65) |
| | β-CH2 | 27.9 | 1.79, m |
| | γ-CH2 | 23.1 | 1.69, m |
| | δ-CH2 | 38.6 | 2.88, m |
| | C=O | 173.6 | - |
| Glycine (1) | α-CH2 | 42.1 | 3.90, 3.86 |
| | C=O | 170.6 | - |
| Serine | α-CH | 52.3 | 4.8, t (5.21) |
| | β-CH2 | 60.3 | 3.59, m |
| | C=O | 169.8 | - |
| N-OH-Orn | α-CH | 52.7 | 4.05, m |
| | β-CH2 | 21.9 | 1.4, m |
| | γ-CH2 | 27.5 | 1.56, m |
| | δ-CH | 47.4 | 3.34, m |
| | COOH | 175.7 | - |
| Glycine (2) | CH2 | 42.1 | 3.78, 3.84 |
| | C=O | 170.7 | - |
| L-Ornithine (2) | α-CH | 53.6 | 4.2, dd (5.80, 8.40) |
| | β-CH2 | 27.9 | 1.79, m |
| | γ-CH2 | 23.1 | 1.69, m |
| | δ-CH2 | 38.6 | 2.88, m |

| | | | |
|---|---|---|---|
| | C=O | 173.6 | - |
| D-Ornithine (2) | α-CH | 53.6 | 4.42, dd (5.80, 8.74) |
| | β-CH2 | 27.7 | 1.85, m |
| | γ-CH2 | 23.8 | 1.65, m |
| | δ-CH2 | 38.6 | 2.88, m |
| | C=O | 174.1 | - |
| 2,3-Dihydroxy benzoic acid (2) | C=O | 170 | - |
| | qC | 115.1 | - |
| | C | 146.4 | - |
| | C | 144.3 | - |
| | CH | 119.6 | 6.91, t (8.65) |
| | CH | 119.6 | 6.68, d (8.07) |
| | CH | 119.3 | 7.08, m |

**Table S8.** Advanced Marfey's analysis of cellulochelin B. Related to Figure S17.

| Amino Acid | Analyte $t_r$ (min) | L-Standard $t_r$ (min) | D-Standard $t_r$ (min) |
|---|---|---|---|
| Serine | 5.69 | 5.67 | 5.87 |
| Ornithine | 3.85, 4.28 | 4.24 | 3.85 |

**Table S9**. Top 10 MS2 fragments of cellulochelin A from *Cellulomonas* sp. strain leaf334

| *m/z* | Intensity |
|---|---|
| 1060.509 | 7.9E+03 |
| 696.3315 | 4.7E+03 |
| 251.1034 | 3.9E+03 |
| 810.4099 | 3.6E+03 |
| 446.2363 | 3.5E+03 |
| 115.0868 | 3.4E+03 |
| 1061.509 | 3.0E+03 |
| 365.1823 | 2.8E+03 |
| 560.3154 | 2.2E+03 |
| 811.413 | 2.2E+03 |

**Table S10**. Top 10 MS2 fragments of cellulochelin B from *Cellulomonas* sp. strain leaf334

| *m/z* | Intensity |
|---|---|
| 714.3406 | 1.0E+04 |
| 464.2461 | 7.8E+03 |
| 251.103 | 7.2E+03 |
| 115.087 | 5.8E+03 |
| 1078.515 | 5.5E+03 |
| 828.4214 | 3.9E+03 |
| 715.3421 | 3.7E+03 |
| 365.1813 | 3.3E+03 |
| 715.3495 | 3.0E+03 |
| 1079.513 | 3.0E+03 |

**Table S11**. List of strains used in the study.

| Strain | Reference |
|---|---|
| *Methylophilus* sp. strain 5 | (2) |
| *Burkholderia ambifaria* BAA244 | (3) |
| *Chromobacterium violaceum* CV017 | (4) |
| *Methylorubrum extorquens* PA1 | (5) |
| *Rhodopseudomonas palustris* CGA009 | (6) |
| *Escherichia coli* MG1655 | (7) |
| *Kushneria konosiri* JCM16805 | (8) |
| *Paracoccus denitrificans* PD1222 | (9) |
| *Cellulomonas* sp. strain Leaf334 | (10) |

### REFERENCES


1. Sandy M, Butler A. 2011. Chrysobactin Siderophores Produced by *Dickeya chrysanthemi* EC16. *J Nat Prod* 74:1207–1212.
2. Chistoserdova L. 2010. Genomes of fifty methylotrophs isolated from Lake Washington. DOE Joint Genome Institute.
3. Barelmann I, Meyer J-M, Taraz K, Budzikiewicz H. 1996. Cepaciachelin, A New Catecholate Siderophore From *Burkholderia* (*Pseudomonas*) *cepacia*. Zeitschrift für Naturforschung C 51:627–630.
4. Wang X, Hinshaw KC, Macdonald SJ, Chandler JR. 2016. Draft Genome Sequence of *Chromobacterium violaceum* Strain CV017. *Genome Announc* 4:e00080-16.
5. Marx CJ, Bringel F, Chistoserdova L, Moulin L, Farhan Ul Haque M, Fleischman DE, Gruffaz C, Jourand P, Knief C, Lee M-C, Muller EEL, Nadalig T, Peyraud R, Roselli S, Russ L, Goodwin LA, Ivanova N, Kyrpides N, Lajus A, Land ML, Medigue C, Mikhailova N, Nolan M, Woyke T, Stolyar S, Vorholt JA, Vuilleumier S. 2012. Complete Genome Sequences of Six Strains of the Genus *Methylobacterium*. *J Bacteriol* 194:4746–4748.
6. Baars O, Morel FMM, Zhang X. 2018. The purple non-sulfur bacterium *Rhodopseudomonas palustris* produces novel petrobactin-related siderophores under aerobic and anaerobic conditions. *Environ Microbiol* 20:1667–1676.
7. Edwards JS, Palsson BO. 2000. The *Escherichia coli* MG1655 *in silico* metabolic genotype: Its definition, characteristics, and capabilities. *Proc Natl Acad Sci USA* 97:5528–5533.
8. Yun J-H, Park S-K, Lee J-Y, Jung M-J, Bae J-W. 2017. Kushneria konosiri sp. nov., isolated from the Korean salt-fermented seafood Daemi-jeot. *Int J Syst Evol Microbiol* 67:3576–3582.
9. Bergeron RJ, Dionis JB, Elliott GT, Kline SJ. 1985. Mechanism and stereospecificity of the parabactin-mediated iron-transport system in *Paracoccus denitrificans*. *J Biol Chem* 260:7936–7944.
10. Vogel CM, Potthoff DB, Schäfer M, Barandun N, Vorholt JA. 2021. Protective role of the *Arabidopsis* leaf microbiota against a bacterial pathogen. *Nat Microbiol* 6:1537–1548.
